# Transcriptional activation of Jun and Fos members of the AP-1 complex is a conserved signature of immune aging that contributes to inflammaging

**DOI:** 10.1101/2022.08.15.503978

**Authors:** Emin Onur Karakaslar, Neerja Katiyar, Muneer Hasham, Ahrim Youn, Siddhartha Sharma, Cheng-han Chung, Radu Marches, Ron Korstanje, Jacques Banchereau, Duygu Ucar

**Author notes:** These authors contributed equally to this paper (Emin Onur Karakaslar, Neerja Katiyar).

## Abstract

Diverse mouse strains have different health and life spans, mimicking the diversity among humans. To capture conserved aging signatures, we studied long-lived C57BL/6J and short-lived NZO/HILtJ mouse strains by profiling transcriptomes and epigenomes of immune cells from peripheral blood and the spleen from young and old mice. Transcriptional activation of the AP-1 transcription factor complex, particularly *Fos, Junb*, and *Jun* genes, was the most significant and conserved aging signature across tissues and strains. ATAC-seq data analyses showed that the chromatin around these genes was more accessible with age and there were significantly more binding sites for these TFs with age across all studied tissues, targeting pro-inflammatory molecules including *Il6*. Age-related increases in binding sites of Jun/Fos factors were also conserved in human peripheral blood ATAC-seq data. Single-cell RNA-seq data from the mouse aging cell atlas Tabula Muris Senis showed that the expression of these genes increased with age in B, T, NK cells, and macrophages, with macrophages from old mice expressing these molecules more abundantly than other cells. Functional data showed that upon myeloid cell activation *via* poly(I:C), the levels of c-JUN protein and its binding activity increased more significantly in spleen cells from old mice compared to cells from young mice. In addition, upon activation, old cells produced more IL6 compared to young cells. In sum, we showed that the aging-related transcriptional activation of *Jun/Fos* members of the AP-1 complex is conserved across immune tissues and long- and short-living mouse strains, possibly contributing to increased inflammation with age.

## INTRODUCTION

Age-related changes in the immune system reduce older individuals’ ability to generate protective responses to immunological threats and lead to increases in diseases and infections^(1, 2)^. Increased inflammation with age (i.e., inflammaging) is one of the hallmarks of immune system aging that is conserved across human and mouse as well as across tissues and strains^(3)^, however, the drivers of this aging signature are mostly unknown^(4)^. Human immune aging studies have mostly been limited to blood since it is easy to access and gives an opportunity to study the status of the peripheral immune system with minimal invasiveness. These studies have uncovered significant age-related changes in gene expression levels in whole blood, as well as in blood-derived peripheral mononuclear cells (PBMCs) and sorted immune cells^(5-7)^. Through genomic profiling, we and others have uncovered that pro-inflammatory molecules are activated with age, whereas molecules related to T cell homeostasis and signaling are downregulated^(4, 5, 7-10)^. Although these studies have described significant age-related changes in transcriptional regulatory programs of immune cells, including the activation of pro-inflammatory programs, they have not pinpointed potential upstream regulators of these genomic alterations.

Mouse models are essential in aging research for establishing age-related changes in various tissues, drivers of these changes, and ways to delay or reverse these changes^(11)^. Most aging studies utilize the longest-living laboratory strain C57BL/6J (B6) with a median life span of 901 days for males and 866 days for females^(12)^. However, there is significant diversity among mouse strains in terms of health and life span that could be exploited to uncover signatures of aging conserved among diverse human populations. An example of such a strain is New Zealand Obese (NZO/HILtJ), which resembles human aging in multiple aspects. First, similar to humans, NZO females live longer than males^(13)^, where the median lifespan is 576 days for females and 423 days for males^(12)^. Second, NZO mice develop obesity, and hence are used as models in Type 2 diabetes (T2D), insulin resistance, and obesity^(14)^ research. Obesity is an epidemic in human populations and a factor that accelerates the mechanisms of aging, significantly contributing to unhealthy aging. Thus, NZO and B6 represent ‘unhealthy aging’ and ‘healthy aging’ models, respectively, enabling us to uncover biomarkers of aging that are consistent despite differences in genetic backgrounds and life and health spans.

In these two strains, we characterized the effects of age on the immune system by profiling peripheral blood leukocytes (PBL) and the cells from the spleen, the largest lymphoid organ, both of which harbor post-differentiated immune cells affected by aging. We also profiled naive and memory CD8^+^ T cells sorted from the spleen, given the importance of CD8^+^ T cells in human aging^(5, 7)^. In these strains and tissues, we studied the effects of age on the transcriptome (*via* RNA-seq), epigenome (*via* ATAC-seq), and cell compositions (*via* flow cytometry) by generating and comparing data from young (3 months) and old (18 months) animals. Together, our data uncovered transcriptional activation of c-Jun and c-Fos members of the AP-1 complex with age as the most conserved biomarker of immune aging in long- and short-living mouse strains and across spleen and blood immune cells. Our results, including functional data on c-JUN binding and IL6 production, suggest that AP-1 complex activation may be a driver of inflammaging. Data from these two mouse strains and human PBMCs can be queried at: http://immune-aging.jax.org.

## RESULTS

### Profiling blood and spleen immune cells of young and old long-lived B6 and short-lived NZO mice

To comprehensively study the effects of aging on the mouse immune system, we generated flow cytometry, RNA-seq, and ATAC-seq data from circulating immune cells (PBLs) and from the spleen in B6 and NZO strains. These strains were selected as representatives of each end of the longevity spectrum of lab mice to mimic the heterogeneity in individuals’ life spans and to uncover conserved immune aging signatures in long and short-living animals. PBL and spleen samples were collected from young (3 months) and old (18 months) mice, which also allows direct age comparison to existing single-cell mouse transcriptomic data from the Tabula Muris Senis atlas^(15)^ (**Figure 1A**). We also sorted and profiled epigenomes and transcriptomes of naïve and memory CD8^+^ T cells from the spleen to study whether age-related changes in the human CD8^+^ cells^(7)^ also exist in mice. Samples that passed quality control (QC) were used in downstream analyses: 74 flow cytometry samples (39 B6, 35 NZO), 103 RNA-seq samples (49 B6, 54 NZO), and 90 ATAC-seq samples (38 B6, 52 NZO) (summarized in **Figure 1A, Table S1**). After QC and filtering, 96,623 consensus ATAC-seq peaks from all cell/tissue types and 18,294 expressed genes from RNA-seq samples were used in downstream analyses. Using flow cytometry, we characterized PBL and spleen cell compositions for a total of 36 cell types including total B (CD19^+)^, CD8^+^ and CD4^+^ T cells and their subsets (**Table S2**).

**Figure 1.**
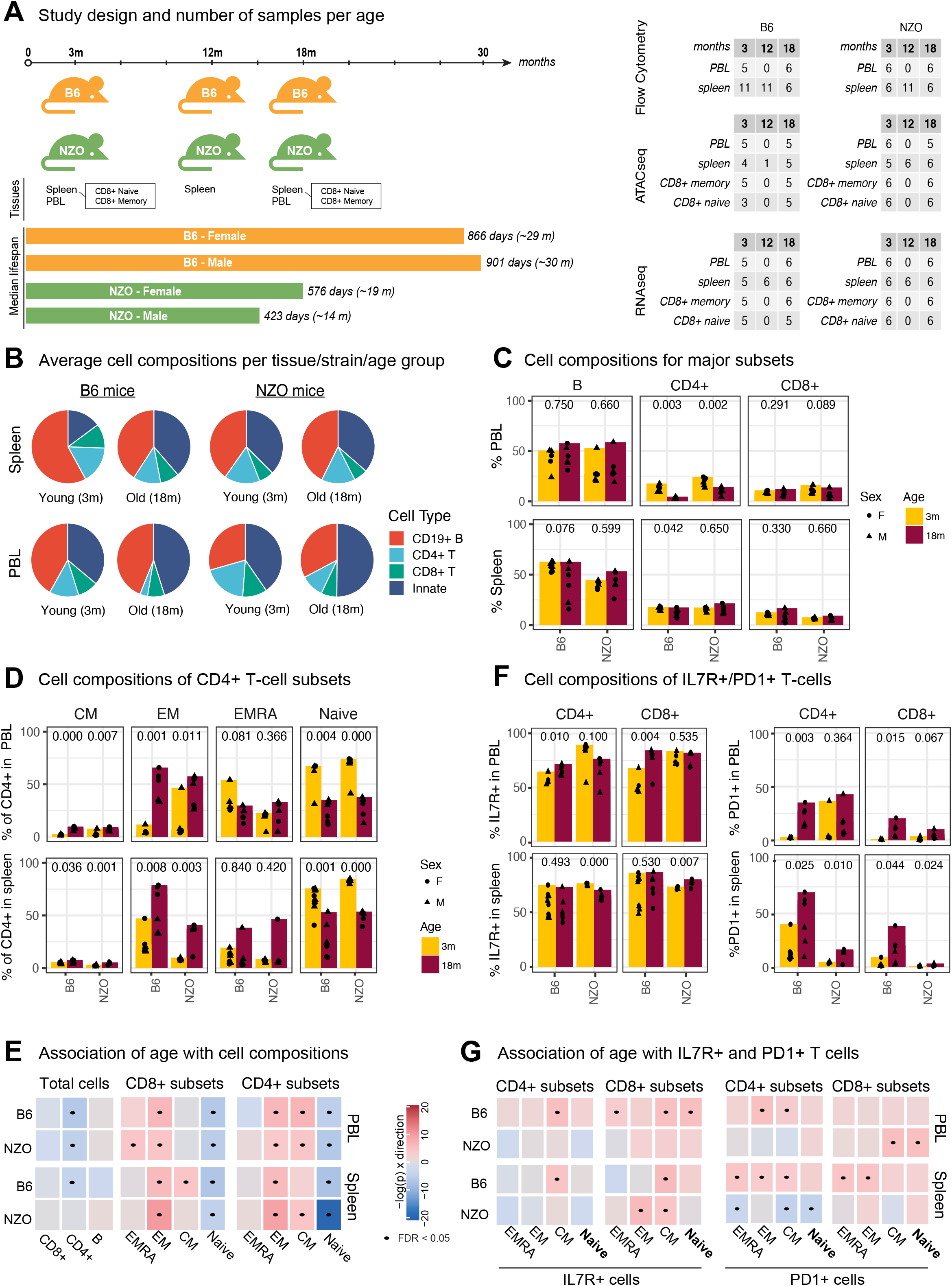
**A** Schematic of study design. PBL and spleen were collected from 3 and 18-month-old mice, and CD8+ T-cells were sorted from spleen. We also collected and profiled spleen from 12-month-old mice. From these samples, ATAC-seq, RNA-seq and flow cytometry data were generated. Summary table of samples, i.e., individual animals: 74 flow cytometry samples (39 B6, 35 NZO), 103 RNA-seq samples (49 B6, 54 NZO), and 90 ATAC-seq samples (38 B6, 52 NZO). **B** Overall cell composition pie charts for PBL and spleen from young and old mice for B, CD4^+^ T, and CD8^+^ T cells. The remaining cells were designated as “innate”. **C** Changes in the cellular composition of major cell types in mouse spleen and PBL with age (x-axis represents age in months, and each dot represents an animal). The jitter around points is for plotting purposes to avoid overlapping dots. **D** Similar to (**C**) for CD4+ T cell subsets (CM: Central Memory, EM: Effector Memory, EMRA: Effector Memory re-expressing *CD45RA*). **E** Summary for the association of age with a given cell type using linear models. Each cell type was colored according to the slope of change per unit of age (in months) and the direction of the slope - red for positive (increase with age) and blue for negative (decrease with age) slopes, with darker colors indicating steeper slope (i.e., more significant changes). Significant associations (FDR 5%) are marked with a dot. **F** Cell composition changes of IL7R^+^ and PD1^+^ T cell subsets. **G** Linear models that associate IL7R^+^ and PD1^+^ T cell percentages to age similar to **E**. p-values in C-D-F are calculated using unpaired t-test.

Principal component analysis (PCA) revealed that RNA-seq, ATAC-seq, and flow cytometry samples were first separated by tissue/cell type and then by strain as expected (**Figure S1A-C**). We quantified how much variation is attributable to meta-data in each modality using principal variance component analysis (PVCA)^(16)^. Tissue/cell type explained most of the variation in both RNA-seq (∼55%) and ATAC-seq (∼41%) data, followed by strain differences (16% in RNA-seq, 19% in ATAC-seq) (**Figure S1D**). Age explained 20% of the variation in the flow cytometry data, due to significant remodeling of PBL and spleen cell compositions with age, whereas it explained around 5% of the variation in RNA-seq and ATAC-seq data. Only <2% of the variation in each modality was attributable to biological sex (**Figure S1D**). These data and analyses suggest that cell type and strain are the main drivers of variation in mouse genomics data, followed by the age of the animals.

### Naive T cell decline is the most significant and conserved age-related cell compositional change

Using flow cytometry, we first, quantified the proportion of CD19^+^ B, CD4^+,^ and CD8^+^ T cells within the spleen and PBL (**Table S2**). The myeloid compartment (granulocytes, monocytes, and dendritic cells - DCs) and NK cells constitute the remainder of the cells (henceforth labeled as ‘innate’ cells). The T cell subsets were further stratified into effector memory (EM, CD44^high^ CD62L^-)^, effector (EMRA, CD44^low^ CD62L^-)^, central memory (CM, CD44^high^ CD62L^+)^, and naive (CD44^low^ CD62L^+)^ cells (**Figure S2A**). Given their importance in T cell homeostasis, signaling^(10, 17)^, and in human aging^(7)^, we also quantified IL7R^+^ and PD1^+^ cells^(4, 18)^.

The majority of mouse spleen cells (45%) and PBLs (37%) were composed of B cells compared to only 5-10% of human PBMCs^(19)^ (**Figure 1B**). Total B and CD8^+^ T cell percentages did not change significantly with age, whereas CD4^+^ T cells declined significantly in PBL (**Figure 1C**). Regarding T cell subsets, the most significant change was the decline of naïve T cells, observed in both B6 and NZO, in both tissues (PBL, spleen), and in both CD4^+^ and CD8^+^ compartments (**Figure 1D, S3A**). We also detected significant increases in EM populations in both compartments. Regression models that associate age with the percentage of each cell type confirmed that declines in naïve CD4^+^ and CD8^+^ T cells and increases in EM CD4^+^ and CD8^+^ T cells were the most significant and conserved age-related changes in cell composition (**Figure 1E**). Interestingly, these cell-compositional changes were similar between the two tissues (PBL, spleen) and between the two strains despite the significant difference in their life spans. Previous mouse studies similarly reported declines in naïve T cells and increases in memory T cells both in spleen^(20)^ and blood^(21)^. Naïve T cell decline with age partially stems from insufficient homeostatic proliferation due to thymus shrinkage and continuous activation of naïve T cells^(22)^; in human PBMCs this decline is more striking for naïve CD8^+^ T cells^(22)^ (**Figure S3B**) whereas in these two mouse strains both CD4^+^ and CD8^+^ T cell compartments were similarly affected with age.

The majority of CD4^+^ and CD8^+^ T cells were IL7R^+^ in both tissues and strains, however, the age-related decline in IL7R^+^ CD8^+^ T cells with age in human PBMCs^(7)^ was not detected in mice (**Figure 1F, S3A**). In contrast, there was a slight increase in IL7R^+^ cells in naïve and memory T cell subsets, particularly for the CD8 compartment of PBLs in B6 mice (**Figure 1G, S3C**). On the other hand, percentages of PD1^+^ T cells increased with age in T cell subsets (**Figure 1F, G**). The increases in PD1^+^ cell percentages were more significant in long-lived B6 compared to short-lived NZO at 18 months of age (**Figure 1G, S3C**), suggesting that the increases in PD1^+^ T cell percentages might not strictly relate to the longevity of the organism. Together, these data and analyses showed that PBL and spleen cell compositions significantly change with age and most of these changes are conserved between the two mouse strains, where declines in naive T cells and increases in EM cells are the most significant and conserved age-related changes.

### Transcriptional activation of Jun/Fos members of the AP-1 complex is a conserved aging signature

We conducted differential gene expression analyses between old and young animals for all studied cells/tissues: spleen, PBL, and naïve and memory CD8^+^ T cells (**Table S3**, FDR = 0.05 and |logFC| > 1). In the spleen tissue, 1960 genes were differentially expressed (DE) in B6 and 1795 in NZO, 675 of which were shared between strains. In PBL we detected 455 DE genes in B6 and 862 DE genes in NZO, whereas in naïve CD8^+^ T cells, 826 and 1718 genes were DE with age in B6 and NZO, respectively. Functional annotation of DE genes using immune modules^(23)^ and cell-specific gene sets from human PBMC scRNA-seq data showed that genes associated with inflammation and the myeloid lineage were activated with age in both strains in PBL and spleen (**Table S4, Figure S4A**), whereas naïve T cell-related genes were inactivated with age - in alignment with cell compositional changes. The most significantly upregulated genes in memory CD8^+^ T cells included age-associated T cell (T^aa^) markers^(4)^, notably *Gzmk* and *Ifng*, in both strains; and memory CD8^+^ DE genes were enriched in marker genes associated with this cell type (**Figure S4B,C**). One of the most prominent changes observed in human CD8^+^ T cells was the downregulation of IL7R and the IL7 signaling pathway with age^(6)^, which was not detected in mouse CD8+ T cells (**Table S4**).

To uncover the most conserved signatures of immune aging, we first compared age-related gene expression changes between the two strains within each tissue, which revealed a positive and significant correlation suggesting that most age-related changes are conserved (correlation coefficient R=0.44 to 0.61, p < 2.2.e-16 for all tissues) (**Figure 2A**). To uncover genes that are the most significantly and robustly associated with aging, we calculated a novel similarity score based on the magnitude of association of genes (MAG) to a phenotype, which was originally developed to assess the similarity between diseases^(24)^. First, we calculated MAG scores for each gene based on their association with aging across the 4 studied cells/tissues as well as the two strains (**Figure 2B, Table S5**). Interestingly, the top three most consistently and significantly aging-associated genes (all upregulated) were *Fos, Fosb*, and *Jun*, which are members of the Activator Protein-1 (AP-1) complex. Gene Set Enrichment analyses of genes sorted with respect to their MAG score confirmed that AP-1 genes were significantly enriched among the most conserved aging genes (NES = 2.096, p=0.005) and showed that 5 genes contributed the most to this enrichment: *Fos, Fosb, Jun, Junb*, and *Maff* (**Figure 2C**). In alignment, DE genes for each cell/tissue type were also enriched for AP-1 genes (**Figure S4C**). For example, the expression of *Fos* gene increased significantly across all 4 cell types/tissues in both strains: ∼7-fold and ∼5-fold increases in NZO and B6 spleen respectively (**Figure 2D, Table S3**). Overall, activation of Jun/Fos family genes was more significant in shorter-living NZO animals in both tissues. These 5 genes effectively separated age groups from each other across all 4 cell types, confirming their robust and conserved association with aging (**Figure 2E**). To study the timing of age-related changes, we have also collected spleen samples from both strains at 12 months. Interestingly, these data showed that these genes were activated at 12 months, suggesting that some of these changes might start in middle-aged animals (corresponding to 40-50 years in humans) (**Figure 2D**).

**Figure 2.**
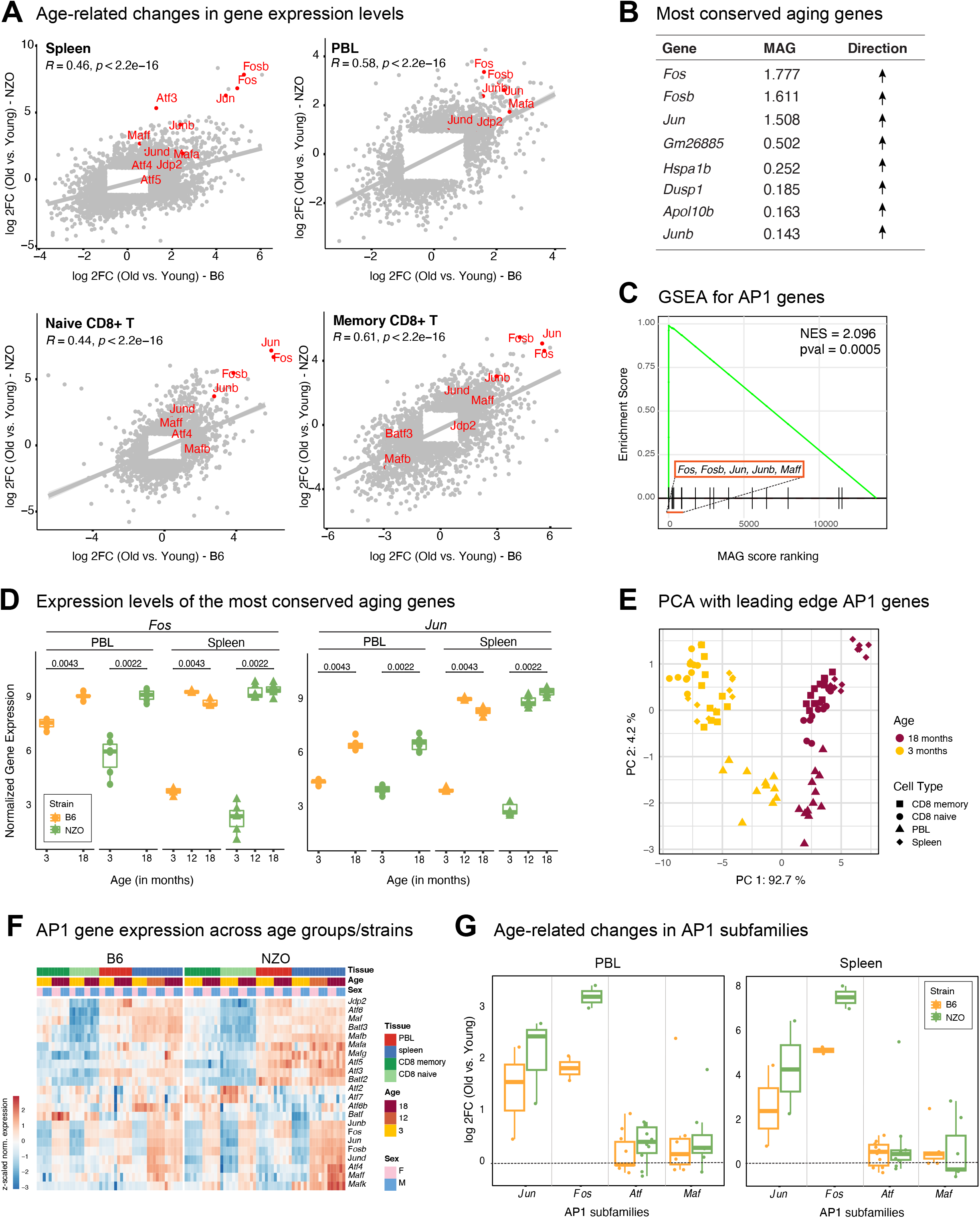
**A** Comparison of fold changes for differentially expressed genes (FDR <0.05) in two strains in each tissue/cell type. Note that transcriptional changes with age are highly conserved across strains. AP-1 complex genes are highlighted in red. **B** Top conserved aging genes (sorted with respect to MAG scores) and the direction of changes with age (upregulated in all cases). **C** Geneset enrichment analysis (GSEA) for MAG-ranked genes using AP-1 complex members as the gene set. Note that top MAG genes are enriched in the AP-1 complex. Leading age genes are denoted. **D** *Fos* and *Jun* expression levels at different ages. Expression values were log(*cpm*) normalized, p-values are calculated using Wilcoxon rank sum test. **E** PCA plot using the expression of leading edge genes (*Fos, Fosb, Jun, Junb, Maff*), note that these five genes clearly separate old samples (dark red) from young ones (yellow) across tissues and strains. **F** Gene expression levels of AP-1 complex genes across all samples. Libraries were normalized using *cpm* function from edgeR package and the gene expression values were z-transformed. **G** Log2 fold changes (positive values refer to upregulation with age) of AP-1 subunits in PBL (left) and spleen (right). Every dot maps to a gene in the corresponding AP-1 subunit. Note that Jun/Fos members are the most strongly affected with aging, all of which are significantly upregulated in both B6 and NZO.

AP-1 is a protein complex that regulates transcriptional responses to diverse stimuli including stress and infections^(25-27)^. Structurally AP-1 is a heterodimer composed of proteins that belong to different subfamilies including Fos (Fos, FosB, Fra-1, Fra-2), Jun (Jun, JunB, and JunD), and ATF (ATFa, ATF-2, and ATF-3)^(281, 29)^. Closer analyses of distinct protein families in the AP-1 complex showed that genes in the JUN/FOS subfamilies were specifically and most significantly activated with age, whereas members of ATF and MAF subfamilies were mostly not associated with age (**Figure 2F, G, Fig S4D, Table S3, and S6**). Together, these data show that the activation of Jun/Fos members in the AP-1 complex is a highly conserved aging signature detected in both strains (B6, NZO) and tissues (PBL, spleen), across all four cell/tissue types, including T cell subsets (naïve and memory CD8^+)^.

### Chromatin accessibility at the Jun/Fos promoters and at their binding sites increases with age

We uncovered differentially accessible (DA) ATAC-seq peaks with age and mapped these to the closest transcription start site (TSS) (**Table S7**, FDR = 0.05 and |logFC| > 1). As expected, genes associated with opening peaks overlapped significantly with genes upregulated with age, whereas closing peak genes overlapped significantly with genes downregulated with age (**Figure S5A)**. Functional enrichment of DA peaks was also in alignment with differential expression results, where the most consistent and strong age-associated signal was the epigenetic activation of pro-inflammatory molecules (**Figure S5B**). We detected significant increases in chromatin accessibility levels around the promoters of Jun/Fos family genes in older samples; this age-related epigenetic activation of Jun/Fos genes was statistically significant and was conserved across strains, tissues, and cell types (**Figure 3A, S5C** for spleen, **S6** for PBL and CD8^+^ subsets).

**Figure 3.**
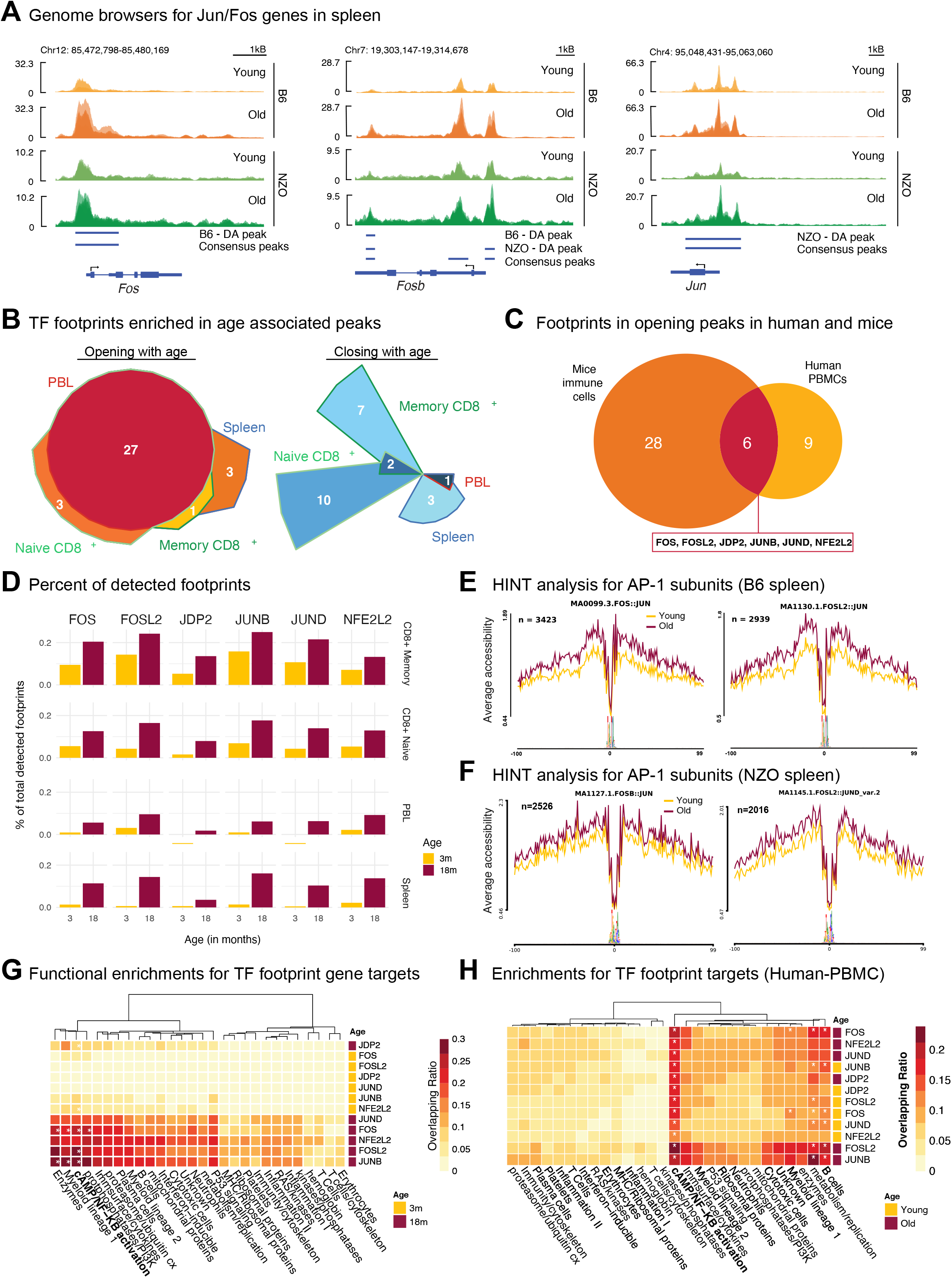
**A** Genome browser tracks that show chromatin accessibility data at the *Jun, Fos* and *Fosb* genes in spleen cells Note that in both strains there is increased chromatin accessibility in older spleen cells at these loci. Bars underneath represent consensus peak and differentially accessible (DA) peak loci. **B** Transcription factor (TF) footprinting enrichment analyses in mice. Left: 34 TF footprints were enriched among opening peaks. 27 were shared across tissue and cell types. Right: 20 TF footprints were enriched among closing peaks; these were cell-specific. (Adjusted q < 0.05 using BiFET). **C** Overlap of footprints enriched in opening peaks in human and mice. 6 proteins were common including five AP-1 members (FOS, FOSL2, JDP2, JUNB, JUND), and a co-factor of the complex NFE2L2. **D** Percent of footprints detected for the shared 6 TFs in each age group and cell type. Counts are normalized with respect to the total number of footprints detected. Note that there are more binding events for these TFs with age. **E, F** Chromatin accessibility levels at the JUN/FOS footprints for (**E**) B6 spleen and (**F**) NZO spleen cells. Note that in older spleen cells, these sites are more accessible. The number of footprints used in each analysis is listed at the top left corner. Functional enrichment of gene targets for conserved age-associated TFs using immune modules in B6 PBL (**G**) and human PBMCs (**H**). Values show the percent of overlap between gene targets and module genes; significant enrichments are indicated with an asterisk (FDR < 0.05). Note the increase in NFkB pathway in older samples.

To uncover whether binding events of Jun/Fos TFs are also affected, we conducted footprinting analyses in ATAC-seq data, which *in silico* infers TF binding events by integrating chromatin accessibility patterns with the underlying TF sequence motifs^(30)^. First, we called TF footprints^(31)^ in mouse samples from pooled young and old animals in 4 different tissue/cell types using known TF motifs^(32)^, as well as in human PBMCs from young and old subjects^(7, 9)^. Samples were pooled to increase the depth of sequencing, increasing the number and quality of detected footprints^(33)^. Next, we identified TF footprints enriched in peaks that were activated (i.e., opening) or inactivated (i.e., closing) with age in each tissue/cell type^(33)^. Interestingly, TF footprints enriched in opening peaks were largely shared across the 4 cell types in mice, in contrast to footprints enriched in closing peaks, which were cell/tissue type-specific (**Figure 3B**). Among the footprints that were enriched in opening peaks with age, six were also enriched in opening peaks from human PBMCs (**Figure 3C, Table S8**). Remarkably, this included five AP-1 complex members (FOS, FOSL2, JDP2, JUNB, JUND) as well as NFE2L2 (i.e., NRF2), a protein that interacts with c-JUN and contributes to the regulation of the NLRP3 inflammasome^(34)^. The baseline activated JUN/FOS status in older mice can be an important modulating element of the biological responses of the aging immune cells. Interestingly, the chromatin accessibility levels around the promoters of these TFs increased in men with aging – inferred from PBMC RNA-seq data from our previous study^(7)^; notably older men experience accelerated myeloid activation and ‘inflammaging’ compared to older women (**Figures S7A**). In alignment with the footprinting results, motif enrichment analyses confirmed that opening peaks with age were enriched in motifs of these TFs, particularly JUN/FOS families both in human and mouse immune cells (**Figure S7B, Table S9**).

In addition to the enrichment of these 6 TFs among opening peaks, footprints for these TFs made up a larger proportion of all detected footprints in older samples across the 4 tissues (**Figure 3D, Table S10**). We also compared the cleavage profiles from the ATAC-seq libraries in young and old mice, to get insights into chromatin accessibility profiles at the binding sites of JUN/FOS TFs^(30, 35)^. Chromatin around their binding sites was more accessible in cells from old mice compared to cells from young mice, which was observed in tissues (PBL, spleen) (**Figure 3E, S7C**) as well as in CD8^+^ subsets in both strains (**Figure 3F, S7D**). To understand which cellular functions are modulated by the increased ‘binding’ of these TFs, we identified their gene targets based on the distance to TSS. These gene targets included members of the Nf-KB pathway (*Rel, Rela, Nfkbiz*), pro-inflammatory cytokines and chemokines (*Il1b, Il6, Il15, Cxcl10*), genes expressed by activated myeloid (*Cd86, Cd44, Il7r, S100a11*) and lymphoid cells (*Cd44, Cd28*)^(19)^, cytotoxic molecules (*Gzmk, Gzmb, Klrg1*), and plasma cell marker *Cd38* (**Table S11**). Our results suggest that these TFs modulate important immune responses in both innate and adaptive immune cells. These gene targets were functionally enriched in pro-inflammatory immune modules and pathways (e.g., Nf-KB activation) across mouse tissues and strains (**Figure 3G, S7E**). Similarly, in human PBMCs, TF gene targets included pro-inflammatory (*FOSL2, LMNA, CASP8, NFKBIZ*), cytotoxic (*GNLY, PRF1, GZMB*), and activated cell markers (*CD44, IL7R*)^(19)^, and were enriched in pro-inflammatory pathways/modules (**Figure 3H, Table S12**). These results support the previously reported cross-regulation of AP-1 complex and NF-kB pathways in myeloid cells^(36, 37)^ and provide further insight into other pathways and functions potentially regulated by these TFs in both myeloid and lymphoid cells. Together, our findings nominate increased JUN/FOS TF activity as a conserved biomarker of immune system aging involved in regulating pro-inflammatory and effector molecule functions, thereby potentially contributing to inflammaging.

### Expression of *Jun/Fos* increases with age across all immune cell types from the spleen

To uncover whether age-related transcriptional activation of Jun/Fos genes stems from specific cell types, we reanalyzed single-cell RNA-seq data from the Tabula Muris Senis consortium^(38)^ using spleen cells from young (3 months) and old (18 months) B6 mice. We annotated spleen cells as B, T, NK cells, or macrophages using known marker genes (**Figure 4A**). In alignment with our flow cytometry data (**Figure 1B**), the majority of spleen cells were composed of B cells, followed by T cells and innate immune cells (**Figure S8A-B**). Next, we studied the activation of the most conserved aging genes (*Jun/Fos/Fosb*) (**Figure 2B**) in these single cells. The expression of these molecules increased with age across all immune cell types (**Figure 4B-C, S8C**). Their activation with age was observed at both the level of expression and the frequency of expression. For example, older B, T, and macrophage cells expressed *Jun* at significantly higher levels than their younger counterparts. In addition, the percentage of cells that expressed these molecules also increased. For example, ∼78% of older macrophages expressed *Fos*, compared to 43% of young macrophages (**Figure 4D**). Despite ubiquitous activation across all immune cell types, macrophages expressed these molecules more frequently than other cell types both in young and old animals among all subsets (**Figure 4D**), emphasizing the importance of *Jun/Fos/Fosb* for inflammatory responses and macrophage functions.

**Figure 4.**
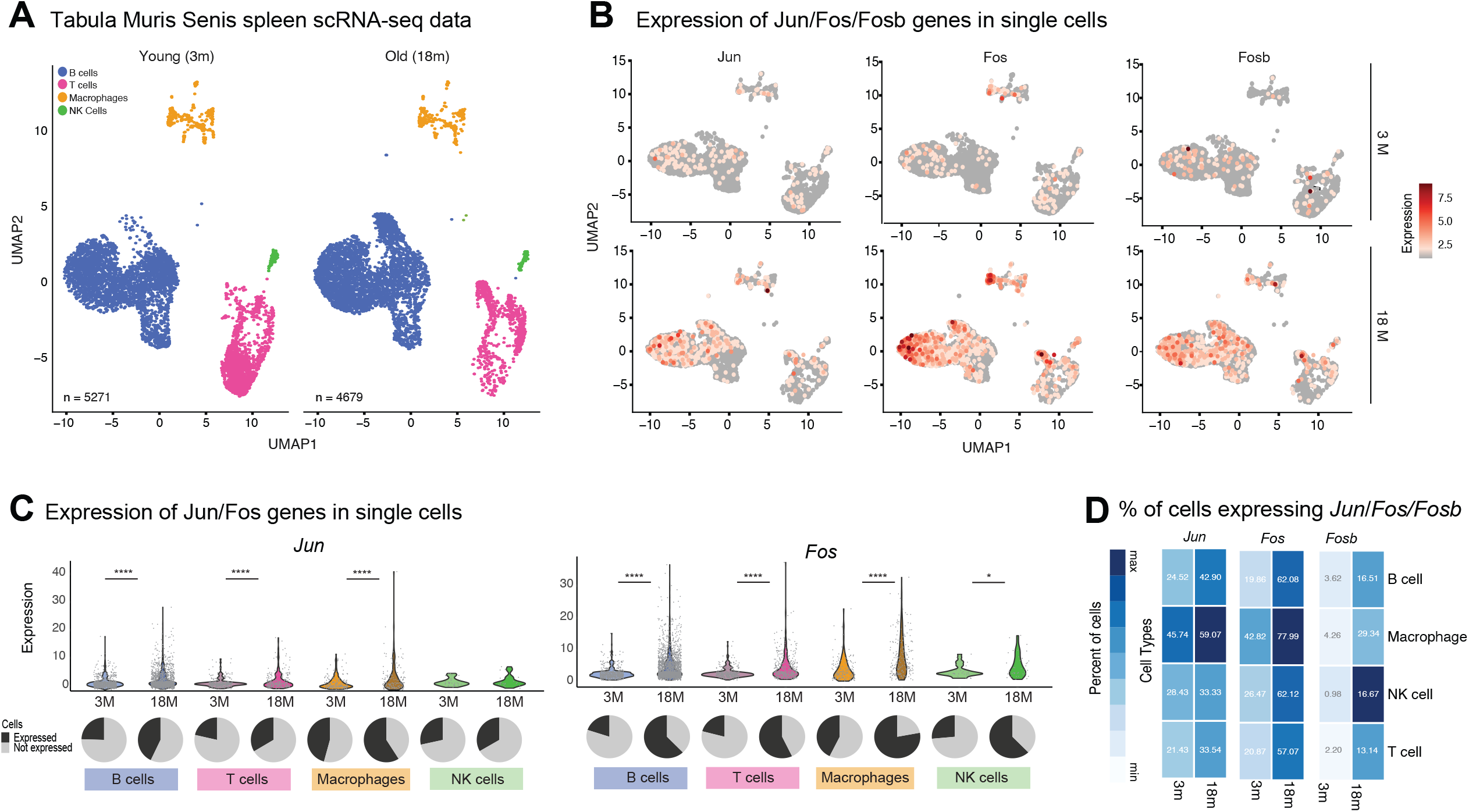
**A** UMAP for the Tabula Muris Senis spleen data using young (3-month) and old (18-month) B6 mice spleen (n=2, per age group). Cell types that have less than 100 cells were removed from the down-stream analyses, resulting in 4 major cell types; B cells, Macrophages, Natural Killer (NK) cells, and T cells (n=9950 cells). **B** The expression levels of the AP-1 complex subunits (*Jun, Fos, Fosb*) in log-normalized scale in young and old cells. Note the increase in expression for all cell types. **C** Violin plots of *Jun* and *Fos* genes in each cell type for young and old mice. Piecharts show the percentage of cells expressing this gene in this cell type. Note that, with age, there is increased expression of these genes both at the level of expression and breadth of expression. **D** Summary for percentage of cells expressing *Jun, Fos, Fosb* genes across cell types in young and old mice.

### c-JUN protein production and binding increase with age upon stimulation *via* Toll-like receptors

An increase in the transcription of genes that form complexes may not necessarily indicate increased function. This is particularly true for the AP-1 protein complex, being a heterodimer composed of c-FOS, c-JUN, and ATF protein families and whose activity depends on the formation of the complex and post-translational modifications such as phosphorylation^(39)^. To understand whether age-related transcriptional/epigenetic activation of *Jun/Fos* affects protein levels and AP-1 function, we studied c-Jun protein expression *via* western blotting and protein binding *via* a c-JUN transcription factor functional assay ELISA kit (**Table S13**). We used splenocytes and not sorted immune cell subsets for these assays since a robust immune response requires cross-talk between different immune cells^(40, 41)^. First, we quantified c-JUN binding activity in nuclear extracts from the B6 spleen cells upon stimulation using 1) anti-CD3/anti-CD28 to stimulate T cells; 2) LPS to stimulate B cells and monocytes *via* TLR4; and 3) poly(I:C) to stimulate monocytes *via* TLR3. The c-JUN binding activity level did not significantly change between age groups upon T cell stimulation, however, it significantly increased with age upon TLR-mediated stimulation (**Figure 5A**), particularly upon poly(I:C) stimulation, which activates monocytes (p=0.0063). TLR3 activation by poly(I:C) has been shown to regulate inflammatory responses in tissues^(42)^. To complement the binding assay, we also quantified the level of c-JUN protein in nuclear and cytosolic fractions of the spleen before and after stimulation by western blotting for c-JUN and loading markers LamininB1 for nuclear extracts and GAPDH for cytosolic extracts. In the nuclear extract, even before the activation of cells, splenocytes from older animals had more c-JUN protein compared to those from younger animals (**Figure 5B**). Upon poly(I:C) stimulation, nuclear and cytosolic c-JUN protein levels increased in both young and old splenocytes as expected, however, the increases were more significant in older cells (**Figure 5B**), in alignment with their transcriptional and epigenetic activation (**Figures 2-3**). Together, our data demonstrate that when activated *via* TLR3 spleen cells from older mice have increased c-JUN protein and c-JUN binding compared to cells from younger mice.

**Figure 5.**
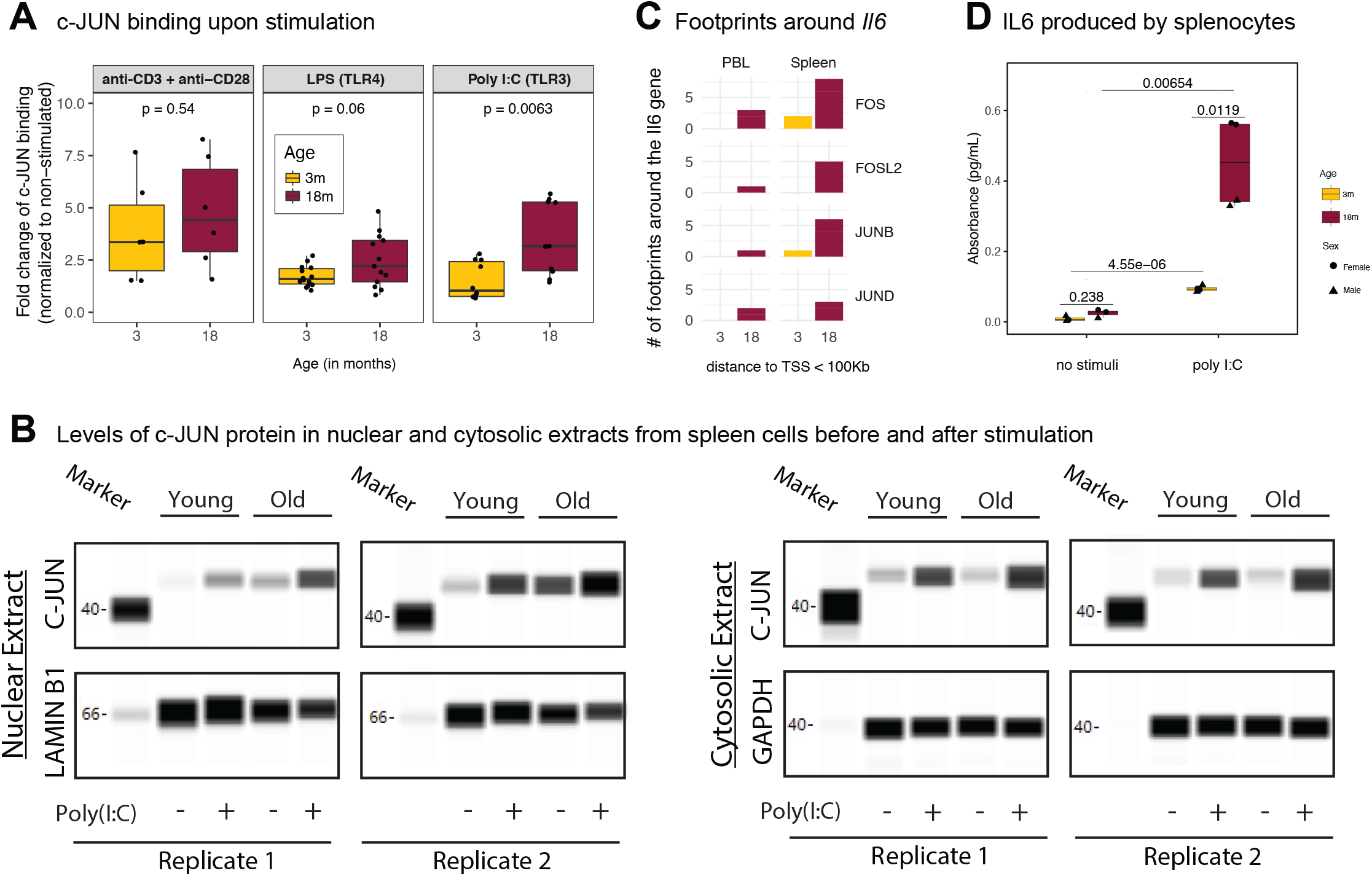
**A** c-JUN binding levels upon different stimuli (anti-CD3+ anti-CD28, LPS and poly(I:C)) for young and old B6 mice spleen. c-JUN binding levels were normalized to the non-stimulated samples. Note the more pronounced increase in c-JUN binding in old spleen cells. P-values are calculated using Wilcoxon rank sum test. **B** Number of footprints detected around the *Il6* gene (100Kb upstream and 100Kb downstream of the TSS from each side) in B6 PBL and spleen for JUN/FOS TFs. Note the increased binding events around this molecule with age. **C** IL6 cytokine levels secreted by splenocytes of young (3 months) and old (18 months) B6 mice spleen. Note the increased production of IL6 with age upon poly(I:C) stimulation. P-values are calculated using unpaired t-test. **D** Western blot of c-JUN proteins from the nuclear and cytosolic extracts of male B6 mice with and without poly(I:C) stimulation. Note that c-JUN levels increase with age in nuclear extracts with age both at baseline and upon poly(I:C) stimulation.

In pooled splenocytes, poly(I:C) stimulation led to the greatest increase in c-JUN binding, which stimulates monocytes *via* TLR3. Monocytes govern the innate immune responses by initiating inflammation, through the production of pro-inflammatory cytokines including IL6. AP-1 complex components are important in regulating inflammation^(43)^, however, their role in inflammaging is unknown. To further explore this connection, we focused on the IL6 cytokine – a canonical marker of inflammaging^(44)^. Footprinting analyses established that footprints for JUN/FOS TFs bound around the *Il6* promoter, and there were more footprints for these TFs around the *Il6* locus with age (**Figure 5C**). To further characterize the downstream effects of increased c-JUN activity with age, we studied whether there is increased IL6 protein production with age in TLR3-activated splenocytes. For this, splenocytes were stimulated *in vitro* with poly(I:C) for 1 day and the supernatant was assayed for IL6. As expected, poly(I:C) stimulation led to increased IL6 levels both in young and old splenocytes, however, this inflammatory response was significantly higher in older cells with age (p= 0.00654) (**Figure 5D**). Interestingly, spleen cells from older females produced significantly more IL6 than spleen cells from older males (p=0.01, **Table S14**). scRNA-seq data from spleen^(38)^ showed that the expression levels of Toll-like receptor genes did not change significantly with age (**Figure S8C**), suggesting that the increased inflammatory responses upon stimulation are modulated by downstream regulators of TLR signaling, not *via* the changes in the receptor expression levels. These data confirm that aged spleen cells are more pro-inflammatory in nature and that increased production and binding of JUN/FOS TFs with age is a potential regulator of these increased inflammatory responses.

## DISCUSSION

To uncover conserved biomarkers and regulators of immune aging, we comprehensively profiled diverse immune cells and tissues in long-living B6 and short-living NZO mouse strains. NZO mice develop T2D and obesity and can thus be considered a model for ‘unhealthy aging’. Despite differences in their health and life spans, our results show significant agreement in age-related changes between the two strains both in flow cytometry (cell compositional) and genomic (cell-intrinsic) data. Using a novel scoring metric, we uncovered that transcriptional activation of JUN/FOS genes of the Activating Protein-1 (AP-1) complex – particularly the upregulation of *Jun, Fos*, and *Fosb* genes – is the most conserved aging signature across the two mouse strains, diverse immune tissues, and cell types. AP-1 is a transcription factor (TF) complex that regulates gene expression programs in response to diverse stimuli, including stress, viral infections^(29)^, and pro-inflammatory signals in concert with the NF-KB pathway^(36, 37, 45)^. ATAC-seq data from the same tissues revealed that there is also epigenetic activation of *Jun/Fos* genes with age. Chromatin accessibility at their promoters increases with age in addition to the chromatin accessibility increases at their binding sites. Western blot data showed that there is more nuclear c-JUN protein in older splenocytes compared to younger ones even at baseline. Furthermore, when myeloid cells within splenocytes are stimulated *via* TLR3 (poly(I:C)), older cells produce more c-JUN and there is more c-JUN binding compared to younger cells. These data suggest that transcriptional activation of *Jun/Fos* genes, whose protein products form the AP-1 complex, is a conserved signature of immune aging that increases the level of c-JUN protein production and binding with age.

Increased inflammation with age (i.e., inflammaging) is one of the hallmarks of aging observed in multiple mouse tissues^(3)^ and also in human cells^(9, 46)^, including the increased levels of IL6 in human serum^(47)^. Several lines of evidence indicate that the activation of *Jun/Fos* genes with age manifests most prominently in the myeloid compartment, specifically monocytes that govern the inflammatory responses. First, poly(I:C) primarily activates monocytes *via* TLR3. TLR3 is a highly conserved molecule that recognizes double-stranded (ds) RNA associated with a viral infection and induces the activity of the interferon response and pro-inflammatory molecules in myeloid cells^(48)^. Second, single-cell RNA-seq data from Tabula Muris Senis^(38)^ showed that, although the gene expression levels of Jun/Fos members increase significantly across all immune cell types with age, older macrophages express these molecules at higher percentages compared to other immune cell types. Footprinting analyses nevertheless suggest that these TFs can target and potentially activate important molecules across distinct immune cell types: pro-inflammatory, cytotoxic, and effector molecules in both myeloid and lymphoid cells. Among the pro-inflammatory molecules, we further studied IL6 as a biomarker of inflammaging. Footprinting analyses demonstrated that JUN/FOS binding events (i.e., footprints) in the vicinity of *Il6* locus increase with age. Furthermore, older spleen cells produce more IL6 upon TLR3-stimulation compared to younger ones. Together with the increased c-JUN protein production and binding in these cells, these results suggest that age-related activation of JUN/FOS TFs with age is a potential upstream regulator of inflammaging.

A prominent age-related change we previously detected in human PBMCs was the chromatin closing and reduced expression of *IL7R* and its downstream molecules in the IL7 signaling pathway^(7)^ that stem from the decline of IL7R^+^ CD8^+^ cells^(7)^. In mouse tissues and strains studied here, we detected neither the downregulation of the *Il7r* gene nor a decline in the percentages of IL7R^+^ T cells. This discordance between human and mouse CD8^+^ T cell aging patterns might be attributable to differences in their antigenic challenges – unlike humans, lab mice live in a highly controlled environment (strict diet, unchallenged immune system). Recent single-cell studies from human and mouse (B6) immune cells have uncovered conserved expansion of GZMK^+^ CD8^+^ T cell populations with age^(4)^. In the present study, we also detected upregulation of marker genes for this T cell population in both strains, confirming that this aging signature is conserved across different strains of mice regardless of their life span.

B6 and NZO strains exhibit differences in terms of the life span and health span of female and male animals, where NZO males live shorter and develop T2D (**Figure 1A**). However, we did not observe significant sex differences in the aging signatures described here, likely because the cohort was not powered to detect sex differences as this was not our main objective. Bigger cohorts will be needed to delve into differences between female and male immune system aging in different mouse strains and their potential implications for human immune aging and vaccine responses. However, we did observe an interesting sex dimorphism in the production of pro-inflammatory IL6 upon stimulation, where female splenocytes from old B6 mice produce significantly more IL6 compared to splenocytes from old males (**Figure 5D**). In alignment with these results, previous studies suggested that testosterone has a suppressing effect on *Jun/Fo*s genes and this plays a role in sex differences observed in human vaccine responses^(49, 50)^. Another human study showed that chromatin accessibility around AP-1 members decreases upon influenza vaccination in blood-derived immune cells – both with trivalent influenza vaccine (TIV) and AS03-adjuvanted H5N1 vaccine in young adults^(51)^. Interestingly, this epigenetic remodeling around the AP-1 complex members boosted responsiveness to vaccines for other viruses – Zika and Dengue^(51)^. Here, we show chronic activation of AP-1 complex members with age. In human PBMCs, AP-1 member gene expression does not significantly increase with age, though increases were more pronounced in men. However, we detected more binding events for JUN/FOS TFs in PBMCs from older adults compared to PBMCs from younger adults, suggesting that the binding activity of these molecules might also be affected with age in human immune cells. Future studies in older adults will be important to uncover the extent of JUN/FOS remodeling with age and the contribution of AP-1 members to reduced vaccine responsiveness in older adults.

## Acknowledgments

We thank Taneli Helenius for aid in scientific writing, members of the JAX Genome Technologies, and Flow Cytometry cores (Heidi Munger, Janet Bakeman, Mary Barter, and Will Schott) for their help with generating the genomic and flow cytometry data. We thank members of the Ucar and Stitzel labs including research interns Emily Lachtara and Lori Kregar, Karolina Palucka, and Nadia Rosenthal for critical feedback during the progress of the study. This study was made possible by the generous financial support of the American Federation for Aging Research (AFAR), the National Institute of General Medical Sciences (NIGMS) under award number GM124922 (to D.U.), and National Institutes of Health (NIH) grants R01 AG052608, R01 AI142086, UH2 AG056925 (to J.B.), P30 AG038070 (The Jackson Laboratory’s Nathan Shock Center of Excellence for the Basic Biology of Aging) (to R.K.), National Cancer Institute P30CA034196 and The PDX R&D Core (The Jackson Laboratory) (to M.H). Opinions, interpretations, conclusions, and recommendations are solely the responsibility of the authors and do not necessarily represent the official views of the National Institutes of Health (NIH).

## Author contributions

DU designed the study with help from JB, MH, and RK. OK, NK, AY, analyzed the data. RK, MH, RM collected samples and generated data. MH, RM, and CC conducted functional experiments. DU, OK, MH wrote the paper. SS developed the R Shiny app. All authors read and edited the manuscript.

## Competing interests

The authors declare no competing financial interests.

While this work was performed and the manuscript was being prepared, J.B. served on the Board of Directors (BOD) for Neovacs; served on the Scientific Advisory Board (SAB) for Georgiamune LLC; BOD member and a stockholder for Ascend Biopharmaceuticals; Scientific Advisory Board (SAB) member and a stockholder for Cue Biopharma; and a stockholder for Sanofi.

JAX (J.B.) and Sanofi entered into a collaborative research agreement to work on a long-read sequencing project that is not related (ended in Jul 2021). Since Aug 2021, J.B. joined Immunai in New York as Chief Scientific Officer (CSO) and continued a limited affiliation with JAX until the end of Feb 2022.

## Data availability

Raw fastq and processed read count files for all samples are deposited to GEO accession code GSE159798.

## Code availability

The code for figures is deposited at https://github.com/UcarLab/mice-aging-project

## METHODS

### Animals and Housing

C57BL/6J (stock 000664) and NZO/HlLtJ (stock 002105) animals were obtained from The Jackson Laboratory and kept in individually ventilated cages with free access to food (5KOG, LabDiet) and water. The pathogen-free room (health status report attached) was kept between 20°C and 22°C, with a 12-hour light:dark cycle. Spleen and blood samples were obtained immediately after euthanasia through cervical dislocation. The mouse study was approved by The Jackson Laboratory’s Institutional Animal Care and Use Committee.

### Flow Cytometry data generation and analyses

Data is obtained from spleen and peripheral blood lymphocytes (PBLs) of C57BL/6J and NZO/HILtJ mouse strains, hereafter B6 and NZO, respectively, at ages 3, 12, and 18 months. Cells were stained with i) CD8 FITC, Clone 53-6.7 BD Bioscience Cat# 553031, used at 1:240 final concentration, ii) CD3e PE, Clone 145-2C11 eBioscience (Now ThermoFisher) cat# 12-0031-85, used at 1:240 final concentration, iii) CD62L PE-Cy7, Clone MEL-14 Tonbo Biosciences Cat# 60-0621-U100, used at 1:480 final concentration, iv) CD44 APC-Cy7, Clone IM7 Tonbo Biosciences Cat# 25-0441-U100, used at 1:240 final concentration and for 30 minutes at 4 degrees C. Propidium Iodide used for cell viability at 0.5ug/ml.

Then cells were sorted on a FACSAria II (BD Biosciences). Briefly, doublets were gated out, Viable PI-cells were gated, CD3e^+^, CD8^+^ cells were gated and subdivided into CD44^Low^, CD62L^High^ Naïve cells and CD44^High^, CD62L^+/-^ Memory cells. Up to 50,000 cells were sorted for ATAC-seq, remaining cells collected for other RNA preparations. Cells were collected in tubes coated with Fetal Bovine Serum (FBS). The percentages of B, CD4, CD8, Naïve CD4, Central memory CD4, Effector memory CD4, Effector memory RA CD4, Naïve CD8, Central memory CD8, Effector memory CD8, Effector memory RA CD8 were measured. In addition, within each of these cell types, the percentages of IL7R^+^ and PD1^+^ cells were measured. After summing up B, CD4^+^ and CD8^+^ T cells, we labelled the rest of the percentages as monocytes since NK cells, neutrophils and other cell types compose only ∼5% of spleen and PBL. Naïve and Memory CD8 T cells were sorted from spleens as follows: Spleens were removed from mice and teased apart in nylon mesh bags in PBS with 2% FBS, 5mM EDTA, and 0.02% Sodium Azide (FACS Buffer). The cells were lysed with “Gey’s Buffer” (a modification of GBSS as described in the attached paper where the Sodium Chloride was exchanged with Ammonium Chloride at the same molar concentration) for 5 minutes and then washed with FACSBuffer and counted to determine concentration. Cells were stained at approximately 10^8/ml with CD8 FITC, CD3e PE, CD62L PE-Cy7, and CD44 APC for 30 minutes at 4 degrees. Cells were washed and resuspended for sorting in FACS Buffer. I sorted Naïve (CD62L^+^, CD44^Low)^ and Memory (CD62L^+/-^, CD44^High)^ CD8^+^, CD3e^+^ cells. To quantify cell compositional changes with age, we built linear models where age is the independent variable and cell type percentage is the dependent variable. For each model, we computed the slope of change per increase in age by 1 unit and a corresponding p-value, which is later corrected using Benjamini-Hochberg procedure.

### RNA Isolation and QC

Total RNA was isolated from 1 million cells using the RNeasy Mini kit (Qiagen), according to the manufacturers’ protocols, including the optional DNase digest step. For samples with fewer than 1 million cells, RNA was isolated using the RNeasy Micro kit (Qiagen). Sample concentration and quality were assessed using the Nanodrop 2000 spectrophotometer (Thermo Scientific) and the RNA 6000 Nano and Pico LabChip assays (Agilent Technologies).

### RNA Library generation

30 ng of total RNA, with the addition of 6 μl ERCC Spike-In Control Mix 1 (Ambion Thermo Fisher) diluted 1:10,000, was used for library construction. Libraries were prepared by the Genome Technologies core facility at The Jackson Laboratory using the KAPA mRNA HyperPrep Kit (KAPA Biosystems), according to the manufacturer’s instructions. Briefly, the protocol entails isolation of polyA containing mRNA using oligo-dT magnetic beads, RNA fragmentation, first and second strand cDNA synthesis, ligation of Illumina-specific adapters containing a unique barcode sequence for each library, and PCR amplification. Libraries were checked for quality and concentration using the D5000 assay on the TapeStation (Agilent Technologies) and quantitative PCR (KAPA Biosystems), according to the manufacturer’s instructions.

### RNA Sequencing

Libraries were pooled and sequenced 75 bp single-end on the HiSeq 4000 (Illumina) using HiSeq 3000/4000 SBS Kit reagents (Illumina), targeting 40 million reads per sample. We obtained RNA-seq data from spleen, PBL and sorted T cells (derived from spleen) of B6 and NZO mouse strains at age 3, 12, and 18 months. Single-end RNA-seq reads were aligned to the mouse genome (mm10) with Bowtie 2^(52)^ and counts were generated with *RSEM*^(53)^. To normalize the raw counts count-per-million (*cpm*) function from edgeR package is used and the genes that are log(cpm)<1 and expressed less than 2 samples were excluded from rest of the analyses. For differential analysis pipeline, however, raw counts are used with the default options of edgeR package^(54)^, and via TMM normalization.

### ATAC-seq library generation

ATAC-seq libraries were prepared using 50,000 cells, as previously described^(55)^, with the following modifications: digitonin was added to the transposition reaction at a final concentration of 0.01%; the transposition reaction was purified using the Genomic DNA Clean & Concentrator-10 kit (Zymo Research Corporation); PCR amplification was carried out using the Nextera DNA Library Prep (Illumina) Index Adapters, Nextera PCR Master Mix, and PCR Primer Cocktail for 10 cycles of PCR; PCR reaction was purified using 1.7x SPRI beads (Agencourt AMPure XP, Beckman Coulter). Libraries were checked for quality and concentration using the DNA High-Sensitivity LabChip assay (Agilent Technologies) and quantitative PCR (KAPA Biosystems), according to the manufacturer’s instructions. Libraries were pooled and sequenced 75 bp paired-end on the NextSeq 500 (Illumina) using NextSeq High Output Kit v2 reagents (Illumina).

### ATAC-seq data analyses

We obtained ATAC-seq data from spleen, PBL and sorted T cells from spleen of B6 and NZO mouse strains at age 3, 12, and 18 months. Paired-end ATAC-seq reads were quality trimmed using *Trimmomatic*^(56)^ and trimmed reads were aligned to mouse genome (mm10) using *BWA*^(57)^. After preprocessing and quality filtering, peaks were called on alignments with *MACS2* using the BAMPE option^(58)^. The consensus peakset for PCA and PVCA plots were generated gathering all peaks from all tissue/cell types, whereas for differential accessibility analyses the samples of the same tissue were merged to generate one consensus peak set by using R package *DiffBind*^(59)^. Peaks only present in at least two samples were included in the analysis. Raw read counts were normalized using the *cpm* function via the log option turned on from edgeR package^(54)^.

### Statistical Methods

Principal Variance Component Analysis (PVCA) was used in order to determine the sources of variability in flow cytometry data^(16)^, which combines the strengths of Principal Component Analysis (PCA) and Variance Component Analyses (VCA). So, using PVCA the proportions variances were attributed to each factor. To compare the normalized gene expressions and peak counts across different tissues and cell types, Wilcoxon rank sum test was used.

### Differential Analyses

To identify differentially expressed genes and differentially accessible peaks between age groups, we used the R package *edgeR*^(54)^ was used. It fits a generalized linear model (GLM) that includes age as a continuous independent variable and read counts from either ATAC-seq or RNA-seq as dependent variables to test for the effect of age on read counts. We stratified data by tissue and strain and fit GLM within strata. In addition to the age, we included sex as a covariate, which did not yield any statistically significant results. P-values for the age effect were adjusted using the Benjamini-Hochberg procedure, and genes or peaks with FDR-adjusted p value < 0.05 were considered differential.

To uncover genes that are the most significantly and robustly associated with aging, we calculated a similarity score based on the magnitude of association of genes (MAG ^(24^)) to a phenotype. The MAG score was calculated by computing the geometric mean of the inverse of ranks of a gene for the 2 strains. The genes were then ranked based on the summation of the MAG score for each gene across the 4 studied cells/tissues. This ranked gene list was provided as input to the gene set enrichment analysis program using AP-1 genes and age-associated T cell markers as gene sets.

### Enrichment Analyses

Immune modules were obtained from human PBMCs^(23)^. Human and mouse orthologs were identified using R package *biomaRt*^(60)^. Consensus peaks were annotated using *HOMER* ^(61)^ and gene-based analyses were restricted to promoter peaks annotated to the nearest transcription start sites (TSS) of expressed genes. Hypergeometric p-value is calculated for each module of the inflammation genesets. Then, p-values are adjusted for multiple hypothesis testing using Benjamini-Hochberg correction. For all analyses, modules that have FDR-adjusted p value < 0.05 considered as enriched.

### Single-cell data Analyses

We have downloaded 10x single-cell RNAseq spleen data of Tabula Muris Senis^(38)^ from USCS browser (https://cells.ucsc.edu/?ds=tabula-muris-senis+droplet+spleen) and transferred it into R environment (v4.0.5). Next, we selected 3- and 18-months samples (2 samples per age point) and ran the standard Seurat (v4.0.2) pipeline with the default parameters (log normalization, 10 PCs, and UMAP for dimensionality reduction). Then, we removed the cell types which had less than 100 cells in total along with the doublet cluster, remaining cells from four major cell types (NK, Macrophage, B, and T cells) were used in all single - cell related analyses. We detected the cluster cell types based on their respective marker genes here; B cell (*Blnk, Cd79a, Cd79b*), T cell (*Cd3d, Cd3e, Cd3g*), NK (*Ncr1, Gzma*), Macrophage (*Itgax*). We have used t-test to quantify the differences between the expression of cells in young (3 mo) vs. old (18 mo) mice samples for *Jun* and *Fos*.

### Footprinting Analyses

ATAC-seq data from spleen and PBL were scanned for TF footprints using the PIQ algorithm ^(31)^. This method integrates genome-wide TF motifs (i.e., position weight matrices) with chromatin accessibility profiles to generate a list of potential TF binding sites that are bound by a TF. The method also produces a quality score for each footprint (positive predictive value). Only the TF footprints with positive predictive values > 0.9 are used in downstream enrichment analyses.

Before footprint calling, we merged samples of the same sex, strain, tissue and age group to increase read depth, which improved the quality and detection power of PIQ. In addition to that we used *SAMtools*^(62)^ to randomly downsample aligned reads from each merged data set to 50 M reads to minimize the impact of the high correlation between library depth and footprint positive predictive values. We used a set of motifs available in the JASPAR 2016 database (n= 454 for human and n=189 for mouse)^(63)^. Finally, footprint calls were further filtered to include in analyses only those associated to TFs that are expressed in the PBL or spleen. For each cell type, we applied *BiFET*^(33)^ to identify TFs whose footprints are significantly more detected in opening/closing peaks compared to background peaks (peaks whose chromatin accessibility do not change with age). In each tissue, we selected TFs whose *BiFET* q-values are less than 0.05 for at least two samples. We followed the same protocols for our previously published human chromatin accessibility data ^(9)^ and merged PIQ calls of young (<40 years) and older individuals (>65 years). For AP-1 complex related TF analyses, we selected each subunit from JASPAR annotated files. Then, the locations of these TFs were annotated to their closest TSS using *ChIPseeker* package^(64)^ (v1.27.3) to uncover the most effected sites. Finally, we calculated enrichment scores of these sites using hypergeometric p-value test and our immune signature gene sets. We used HINT-ATAC(65) which is part of the Regulatory Genomics Toolbox (RGT). The consensus peak files for young (3 months) and old mice (18 months) along with respective merged bam files were provided as input to the HINT program. To find TFs associated with a particular cellular condition, we checked for motifs overlapping with predicted footprints to find motif predicted binding sites (MPBS) with JASPAR motifs. Finally, HINT was used to generate average ATAC-seq profiles around binding sites of a TF.

Spleens were removed from the euthanized mice, rinsed twice in DPBS, and ground through 0.45uM filters into 5 mL of media (DMEM (Gibco/Thermo-Fisher, 12430112) with 10% FBS (Hyclone/Thermo-Fisher, SH3007003), HEPES (Thermo-Fisher, 15630080) and Pen/Step/Glutamine (Thermo-Fisher, 10378016)) in 6 well plates. The red blood cells were lysed using Red Blood Cells Lysing solution as per the manufacturer’s instructions (Sigma). The resulting splenocytes were suspended in 1 million per mL (∼ 5mL) and stimulated for 2 hours with antiCD3 and antiCD28 (1ug/mL, for both, 100uL plate coated; kind gift of Dr. D.V. Serreze) or LPS 10ug/mL or Poly I:C (both LMW and HMW;10ug/mL each) (Invitrogen;tlr-kit1mw)

After 2 hours, lysates were extracted from splenocytes, both activated and unactivated, according to the manufacturer’s instruction of Nuclear Extraction Kit (Abcam, ab113474), and quantified by Pierce™ Coomassie (Bradford) Protein Assay Kit (Thermo-Fisher, 23200). The c-Jun transcription factor was assayed by c-Jun Transcription Factor Assay Kit (Colorimetric) according to manufacturer’s instruction. For the cytokine experiments, 10 million cells were left in 0.5mL in activation media overnight (+18 hours), and IL6 amount in the supernatants was measured by Mouse IL-6 ELISA kit (Abcam, ab222503) according to supplier’s protocol.

### Cell stimulation and immunoblotting experiments

Mice spleens were removed aseptically and transferred to a Petri dish where they were minced and filtered through a 40-μm nylon cell strainer containing DMEM medium. The cellular suspension was centrifuged to yield a pellet and then depleted of erythrocytes by resuspending the cells in pre-chilled Red Blood Cell Lysis solution (Sigma). The spleen cells were washed twice in DMEM medium containing 25mM HEPES, 1mM L-glutamine, 1% penicillin/streptomycin, and 10% heat-inactivated FCS. Viability of the spleen cells was greater than 90 %. Five million spleen cells per condition were suspended at 1×106/ml in 10-ml Petri dishes pre-coated with 100 μg anti-CD3/CD28 or in culture medium supplemented or not with either LPS (1 μg/ml) or Poly (I:C) (both high and low molecular weight, 10 μg/mL each) (Invivogen). After 2 hours, cells were lysed, and the nuclear extracts were isolated using Nuclear Extraction Kit (Abcam) according to the manufacturer’s instructions. Protein content of the nuclear extracts was quantified by Pierce™ Coomassie (Bradford) Protein Assay Kit (Thermo-Fisher) and the activation of the c-Jun transcription factor was assayed by c-Jun Transcription Factor Assay Kit (Abcam) according to the manufacturer’s instruction. Secretion of IL-6 was quantified from the supernatant of 10 million spleen cells stimulated as above, after 24 hours using Mouse IL-6 ELISA kit (Abcam).

The capillary immunoblotting analysis was performed, using Wes (ProteinSimple, Santa Clara, CA, USA), according to the ProteinSimple user manual. The lysates of the primary male mice splenocytes were mixed with a master mix (ProteinSimple) to a final concentration of 1 × sample buffer, 1 × fluorescent molecular weight marker, and 40 mM dithiothreitol and then heated at 95 °C for 5 min. The samples, blocking reagents, primary antibodies, HRP-conjugated secondary antibodies, chemiluminescent substrate (ProteinSimple), and separation and stacking matrices were also dispensed to the designated wells in a 25-well plate. After plate loading, the separation electrophoresis and immunodetection steps took place in the capillary system and were fully automated. A capillary immunoblotting analysis was carried out at room temperature, and the instrument’s default settings were used. Capillaries were first filled with a separation matrix followed by a stacking matrix, with about 40 nL of the sample used for loading. During electrophoresis, the proteins were separated by molecular weight through the stacking and separation matrices at 250 volts for 40–50 min and then immobilized on the capillary wall, using proprietary photo-activated capture chemistry. The matrices were then washed out. The capillaries were next incubated with a blocking reagent for 15 min, and the target proteins were immunoprobed with primary antibodies followed by HRP-conjugated secondary antibodies The antibodies of GAPDH (sc-25778, 1:200, Santa Cruz Biotechnology), Lamin B1 (12586S, 1:100, Cell Signaling Technology), c-Jun (9165S, 1:50, Cell Signaling Technology), were diluted in an antibody diluent (ProteinSimple).

## Figure Captions

**Figure S1.**
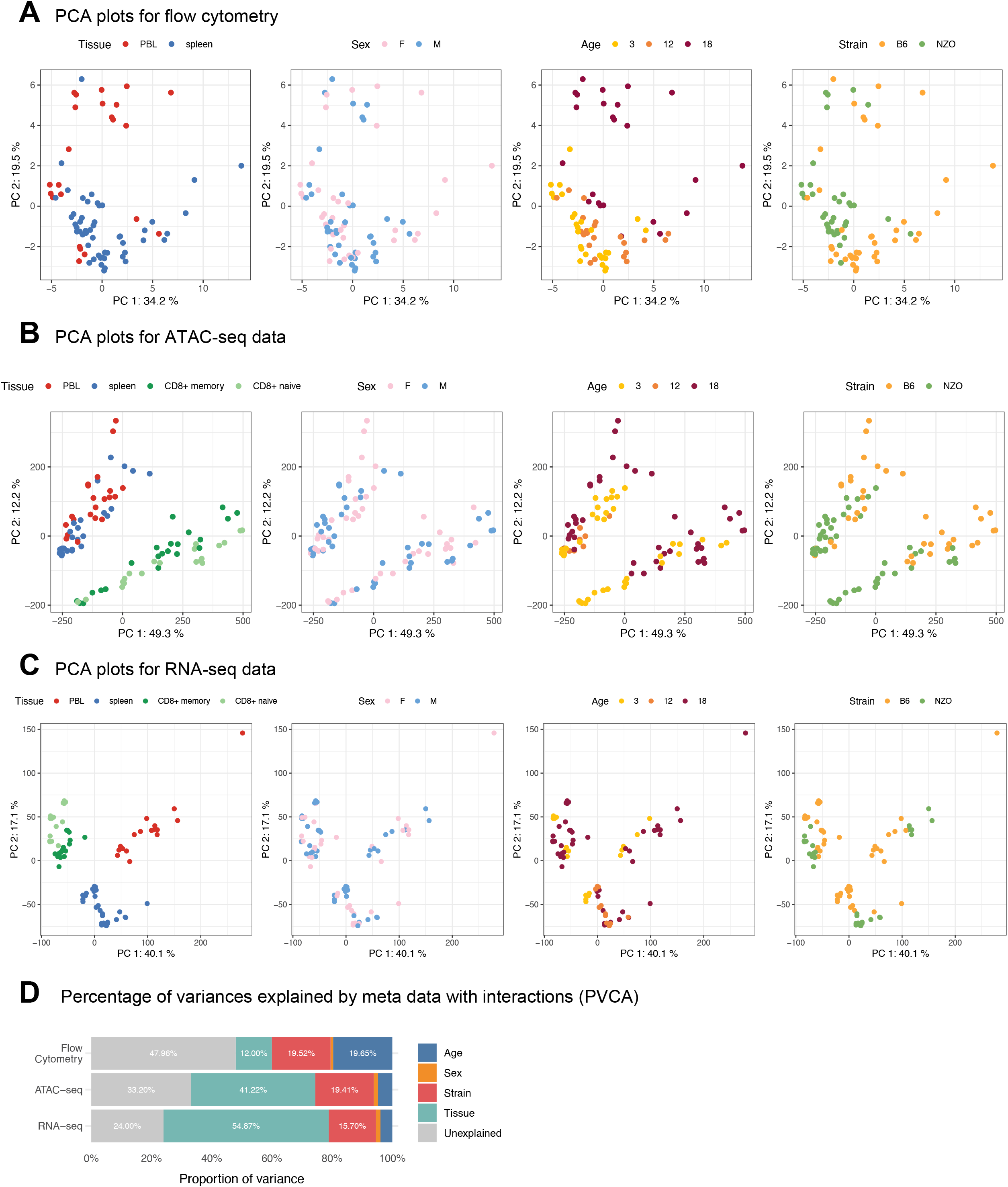
PCA results for (**A**) flow cytometry (n = 33 cell subsets), (**B**) ATAC-seq (n = 96,623 peaks) and (**C**) RNA-seq (n = 18,294 genes). Tissue/cell type information was overlaid onto PC1 and PC2 dimensions and percentages indicate the percent of variation explained with the given principal component. **D** The variances of the datasets were attributed to metadata (sex, age, strain, and tissue/cell type) using principal variance component analyses (PVCA).

**Figure S2.**
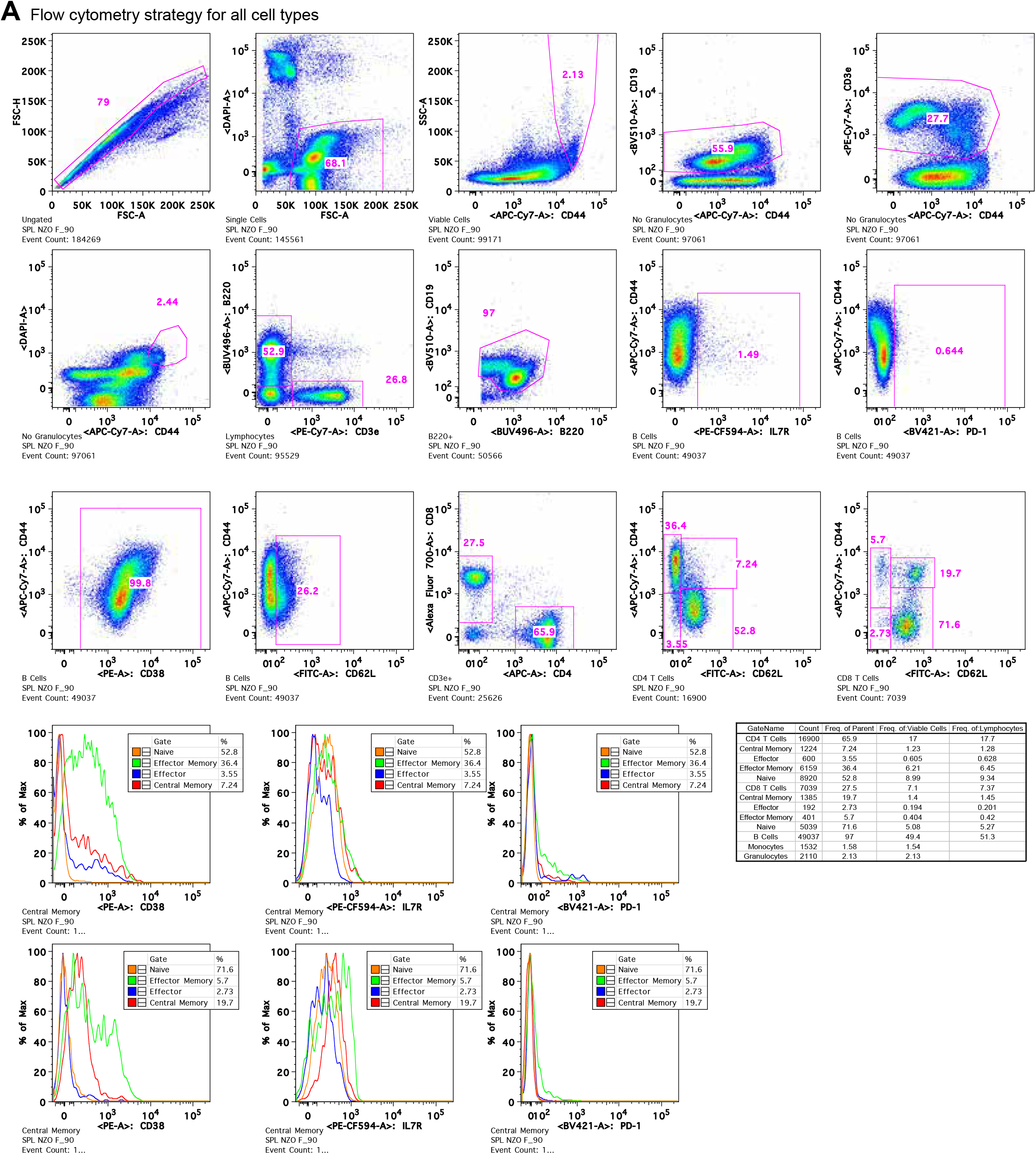
**A** Sample gating strategy for all cell types and subsets. Cells were sorted on a FACSAria II (BD Biosciences). Briefly, doublets were gated out, viable PI-cells were gated, CD3e^+^, CD8^+^ cells were gated and subdivided into CD44^Low^, CD62L^High^ Naïve cells and CD44^High^, CD62L^+/-^ Memory cells. Cells were collected in tubes coated with fetal bovine serum (FBS). The percentages of B, CD4^+^ T, CD8^+^ T, Naïve CD4^+^ T, Central memory CD4^+^ T, Effector memory CD4^+^ T, Effector memory RA CD4^+^ T, Naïve CD8^+^ T, Central memory CD8^+^ T, Effector memory CD8^+^ T, and Effector memory RA CD8^+^ T cells were measured. In addition, within each of these cell types, the percentages of IL7R^+^ and PD1^+^ cells were measured.

**Figure S3.**
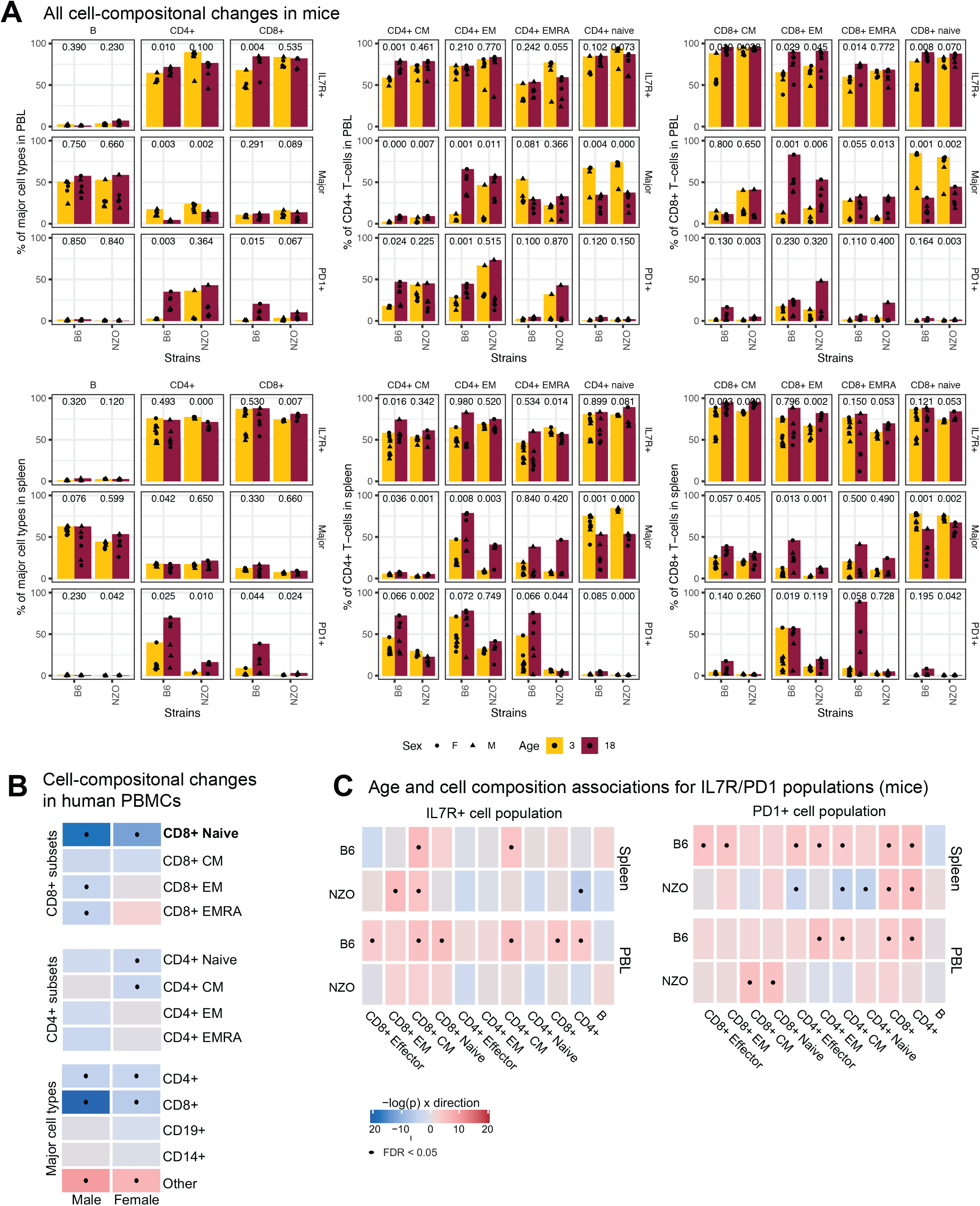
**A** Changes in cellular composition in mouse spleen and PBL with age (x-axis represents age in months, and each dot represents an animal). The jitter around points is for plotting purposes to avoid overlapping dots. P-values are calculated using unpaired t-test **B** Association summary heatmap of human PBMCs for associations of age with indicated cell type using data from previously published human PBMCs^(9)^. Colors (slope) and dots (FDR < 0.05) are as in Figure 1. **C** Association between age and IL7R^+^ and PD1^+^ T cell populations.

**Figure S4.**
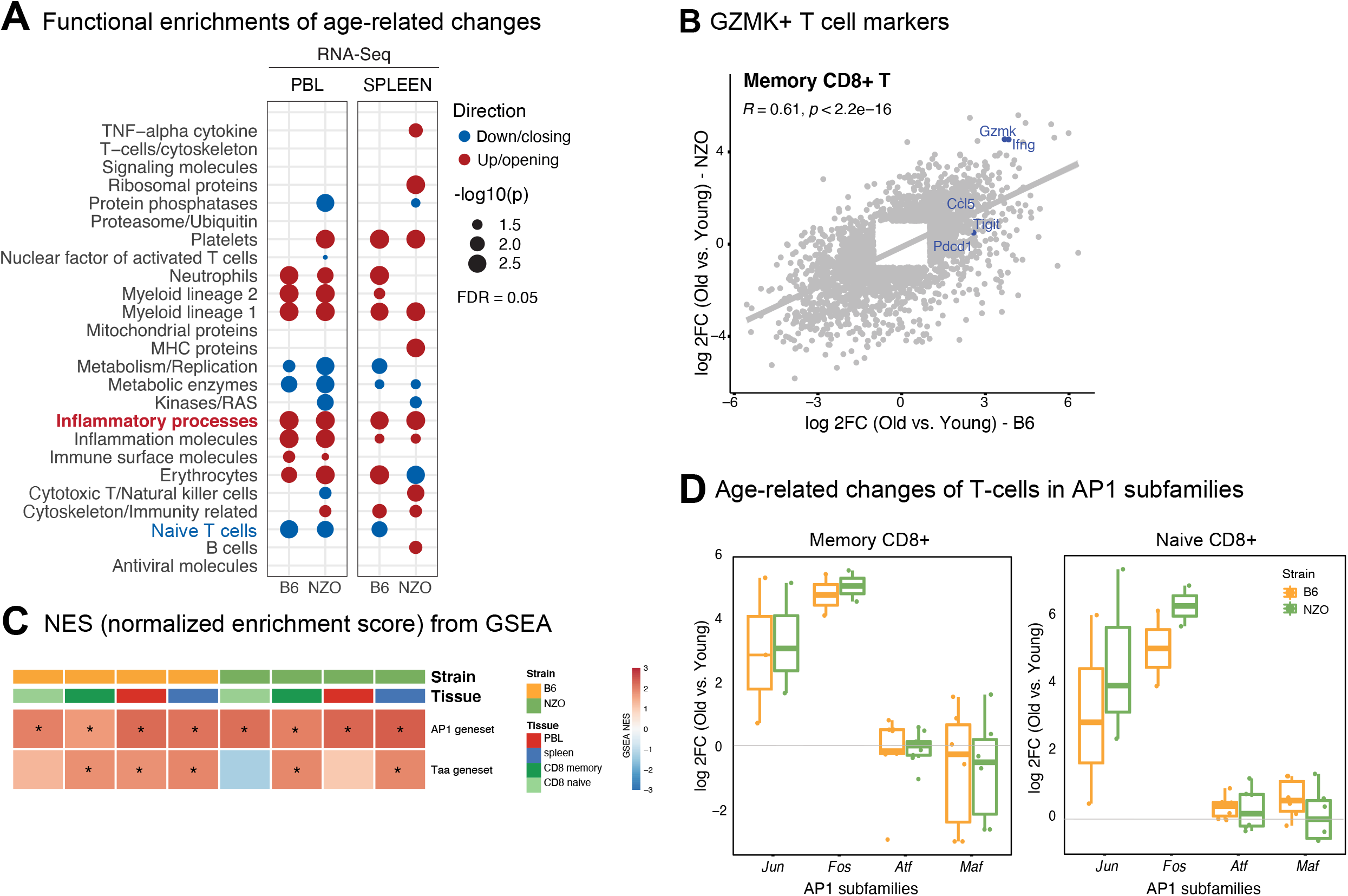
**A** Functional annotations of differential genes using GSEA; p values were calculated using 1000 permutations and corrected using the Benjamini-Hochberg procedure. Only the pathways that have FDR < 0.05 are shown with bubbles. The “Inflammatory processes” module was upregulated/opening across tissue and strain types. **B** Comparison of fold changes for differentially expressed genes (FDR <0.05) in two strains in memory CD8+ T cells, where marker genes for GZMK^+^ CD8^+^ T cells are marked in blue. Note that in both strains these genes are significantly upregulated. **C** Normalized enrichment scores from GSEA analyses using AP1 genes and age-associate T cell marker genes (Taa) as genesets. Note that AP-1 genes are enriched across all tissues. FDR < 0.05 groups were indicated with an asterix. TCT: tissue/cell type **D** Log2 fold changes (positive values refer to upregulation with age) of AP-1 subunits in CD8^+^ subsets.

**Figure S5.**
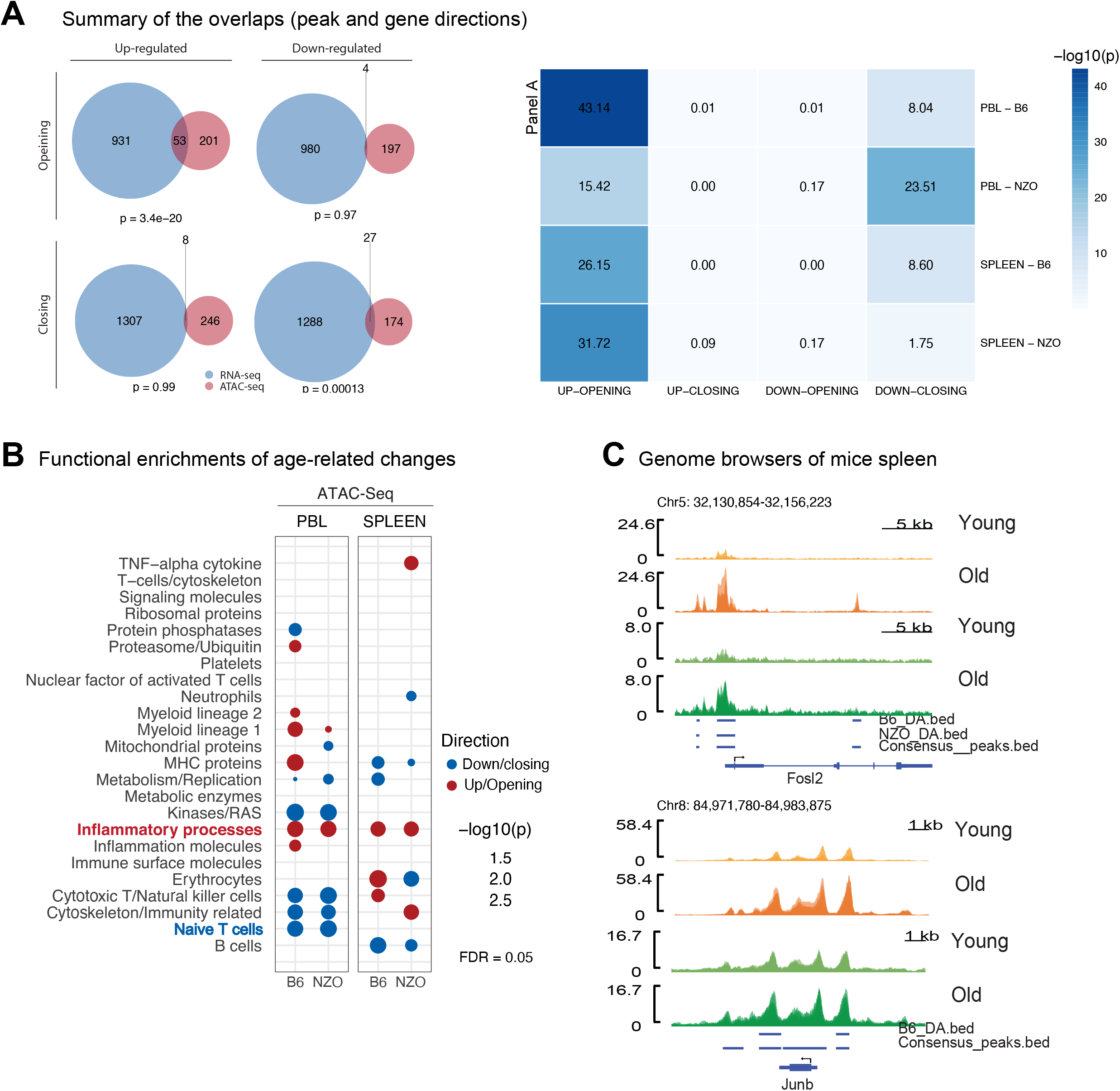
**A** Overlap between RNA-seq and ATAC-seq remodeling with age. As expected, upregulated genes and opening peak genes overlapped significantly, similarly downregulated genes and closing peak genes overlapped significantly. Venn diagrams show overlaps for B6 PBL (left), Heatmap summarizes all comparisons (right). Overlap p-values are calculated using hyper-geometric test. **B** Functional annotations of differentially accessible peaks using GSEA; p values were calculated using 1000 permutations and corrected using the Benjamini-Hochberg procedure. Only the pathways that have FDR < 0.05 are shown with bubbles. The “Inflammatory processes” module was upregulated/opening across tissue and strain types. **C** Genome browser tracks to show chromatin accessibility levels at *Junb* and *Fosl2* genes in spleen. Bars underneath represent consensus peak and differentially accessible (DA) peak loci.

**Figure S6.**
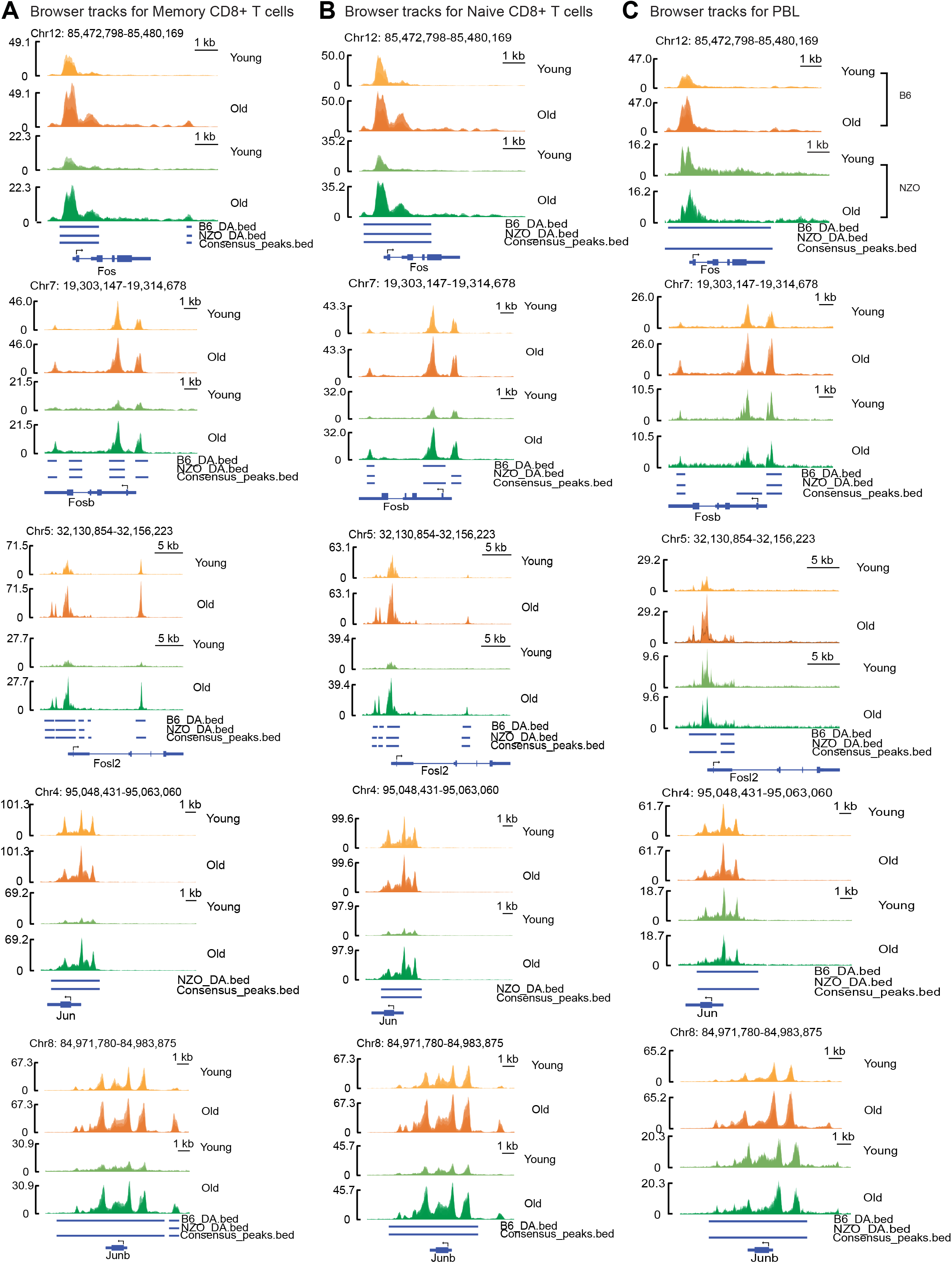
**A, B, C** Genome browser tracks to show chromatin accessibility levels at *Fos, Fosb, Fosl2, Jun, Junb* genes in (**A**) memory CD8+ T cells, (**B**) naïve CD8+ T cells, and (**C**) PBL. Note that the chromatin around these genes are more accessible with age across all studied tissues/cell types. Bars underneath represent consensus peak and differentially accessible (DA) peak loci.

**Figure S7.**
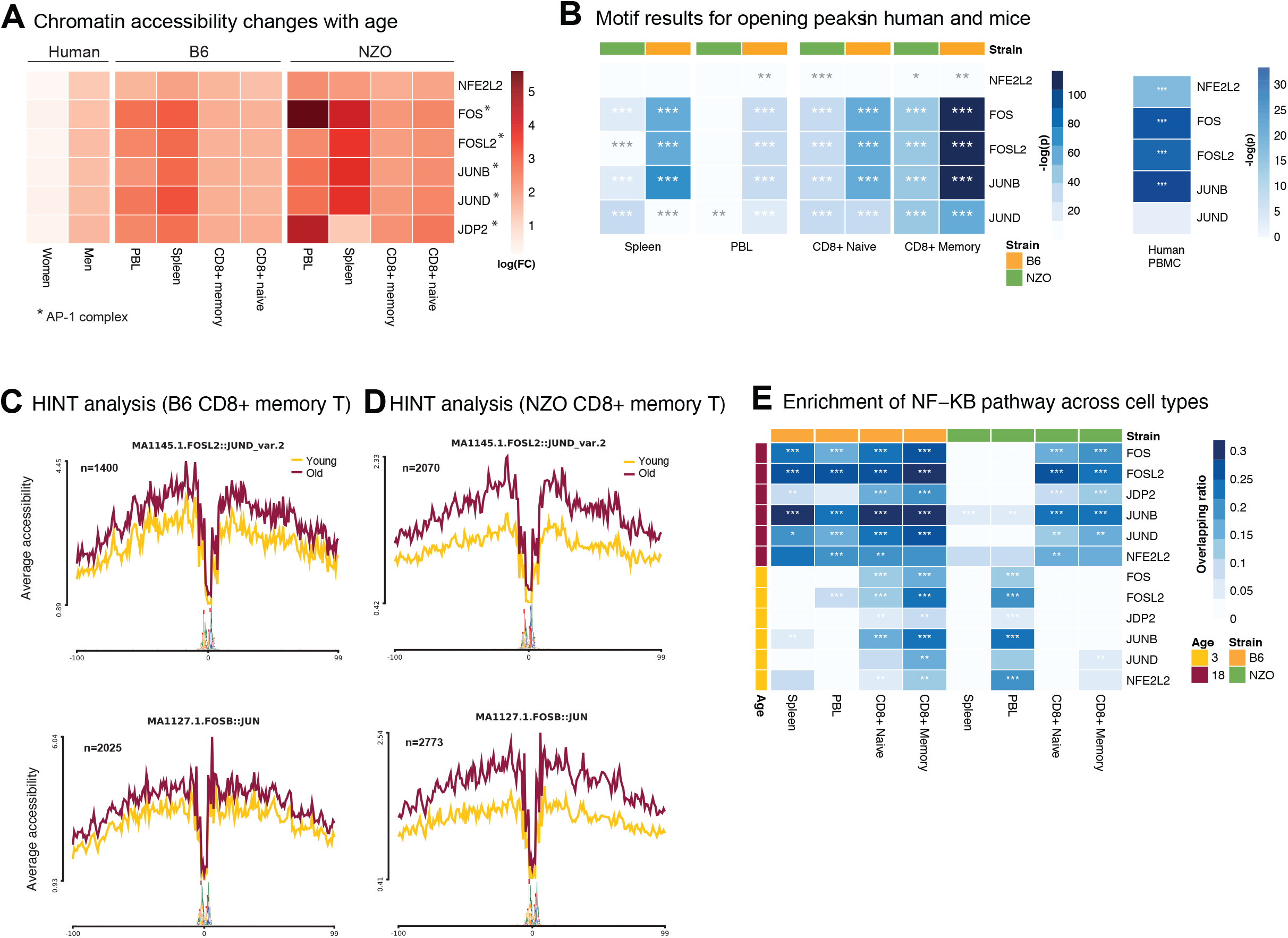
**A** Age-related changes (positive log FC indicates increase with age) in the chromatin accessibility levels of 6 shared TFs. Note the conserved accessibility increase across mice strains and tissues. Activation in human PBMCs is observed in male samples. AP-1 complex members are marked with an asterisk. **B** HOMER motif enrichment results for opening peaks in mice cells/tissues and human PBMCs. **C, D** Chromatin accessibility levels at the JUN/FOS footprints for memory CD8+ T cells in B6 (**C**) and NZO (**D**). **E** Overlap of gene targets for conserved TF footprints with the NFKB pathway genes across cell types and strains. (Enrichment FDR < 0.05*, 0.01**, 0.001***). Note that footprints detected in old samples have higher enrichment and overlap. Overlap p-values are calculated using hyper-geometric test.

**Figure S8.**
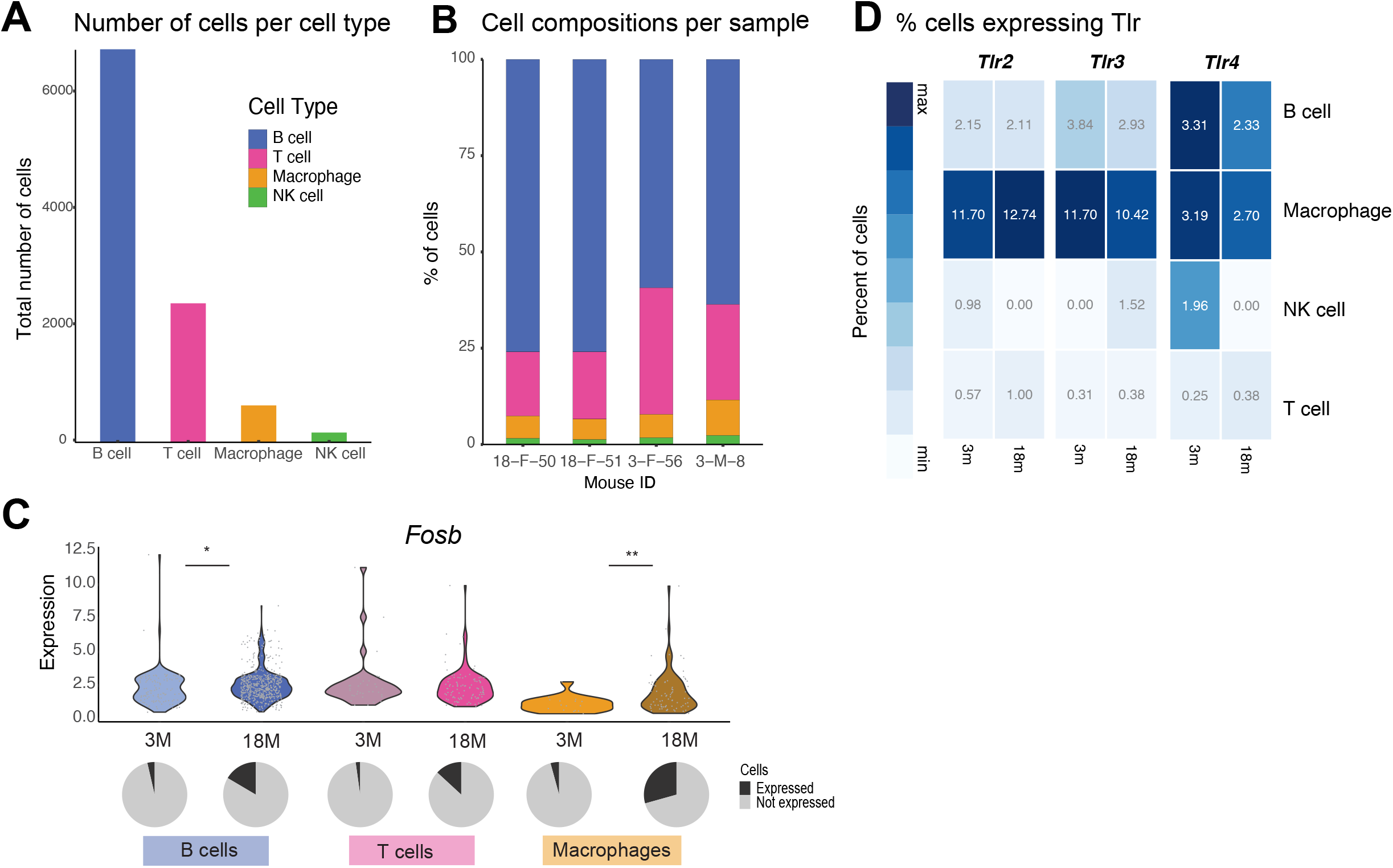
**A** Number of cells per cell type in the Tabula Muris Senis spleen data. Cell types that had less than 100 cells were removed from downstream analyses, resulting in 4 major cell types (B, T, Macrophage and NK cells). **B** Cell type compositions of the four B6 samples. **C** Violin plots of *Fosb* gene in each cell type for young and old mice expect for NK cells, where only a few cells expressed this molecule. Piecharts show the percentage of cells expressing this gene. Note that, with age, there is increased expression of these genes both at the level of expression and breadth of expression. **D** Percentage of cells expressing *Tlr2-4*. Note that TLR expression is not significantly affected with age.

## Supplementary Tables

**Table S1**. RNA-seq and ATAC-seq sample QC

**Table S2**. Flow cytometry data

**Table S3**. Differentially expressed (DE) genes with age

**Table S4**. GSEA enrichments for DE genes

**Table S5**. MAG scores for the genes across cell types/tissues

**Table S6**. AP-1 complex members and families

**Table S7**. Differentially accessible (DA) peaks with age

**Table S8**. BIFET footprint enrichments in opening/closing peaks for mice and human

**Table S9**. HOMER TF motif enrichments in opening/closing peaks

**Table S10**. Footprint counts for aging-associated TFs

**Table S11**. Functional enrichment of TF footprint gene targets (mice)

**Table S12**. Functional enrichment of TF footprint gene targets (human)

**Table S13**. cJUN binding data

**Table S14**. IL6 cytokine data

## Notes

http://immune-aging.jax.org

https://github.com/UcarLab/mice-aging-project

